# Toward high-throughput predictive modeling of protein binding/unbinding kinetics

**DOI:** 10.1101/024513

**Authors:** See Hong Chiu, Lei Xie

## Abstract

One of the unaddressed challenges in drug discovery is that drug potency determined *in vitro* is not a reliable indicator of drug activity in humans. Accumulated evidences suggest that *in vivo* activity is more strongly correlated with the binding/unbinding kinetics than the equilibrium thermodynamics of protein-ligand interactions (PLI). However, existing experimental and computational techniques are insufficient in studying the molecular details of kinetics process of PLI. Consequently, we not only have limited mechanistic understanding of the kinetic process but also lack a practical platform for the high-throughput screening and optimization of drug leads based on their kinetic properties. Here we address this unmet need by integrating energetic and conformational dynamic features derived from molecular modeling with multi-task learning. To test our method, HIV-1 protease is used as a model system. Our integrated model provides us with new insights into the molecular determinants of kinetics of PLI. We find that the coherent coupling of conformational dynamics between protein and ligand may play a critical role in determining the kinetic rate constants of PLI. Furthermore, we demonstrate that the relative movement of normal nodes of amino acids upon ligand binding is an important feature to capture conformational dynamics of the binding/unbinding kinetics. Coupled with the multi-task learning, we can predict combined k_on_ and k_off_ accurately with an accuracy of 74.35%. Thus, it is possible to screen and optimize compounds based on their binding/unbinding kinetics. The further development of such computational tools will bridge one of the critical missing links in drug discovery.

## Significance

Drug efficacy and side effect are often dependent on the life-time rather than the binding affinity of drug-target complex. The ignorance of drug binding/unbinding kinetics seriously hinders the development of efficient and safe therapeutics. For the first time, we integrate physically-based modeling with multi-task learning to investigate the molecular determinants of protein binding kinetics as well as efficiently and accurately predict the kinetic rate constants of drug-target complex. Such computational tools will allow us not only to elucidate novel mechanisms of protein binding/unbinding process but also to screen and optimize compounds based on their kinetic property. This will bridge one of the critical missing links between *in vitro* drug screening and *in vivo* drug efficacy and toxicity.

## Introduction

Target-based and cell-based screenings are the two major approaches in the early stage of drug discovery. In both of these two technologies, one of the unaddressed fundamental challenges is that drug potency measured *in vitro* may not be a reliable indicator of drug efficacy and toxicity in the human body. In the compound screening and lead optimization, equilibrium thermodynamics constants such as half maximal inhibitory concentration (IC_50_) or dissociation constant (K_d_) have been used as the measures of drug potency for years. As molecules in the human body are in a non-equilibrium condition, the activity of a drug depends on, not only how strong it interacts with the protein, but also how easy it hits the target and how long it resides in the target. Increasing body of evidence suggests that drug activity *in vivo* is not defined by equilibrium conditions measured *in vitro,* but rather depends on the residence time (τ = 1/ k_off_) of the receptor-ligand complex *in vivo* in a number of cases [1]. The longer residence time will increase the efficacy of the drug. For example, geldenamycin has low affinity for Heat shock protein (Hsp90) *in vitro* with IC_50_ ~ 1 μM, in comparison to its nanomolar effects *in vivo* [1,2]. Copeland et al. analyzed the results of the experiment of mutation-based resistance to inhibitors of HIV-1 protease, and concluded that the essential factor for sustained drug efficacy *in vivo* is residence time but not binding affinity [3]. Pan et al. reported that residence time is highly correlated with functional efficacy of a series of agonists of the A_2A_ adenosine receptor (r^2^ = 0.95), but there is little correlation with binding affinity (r^2^ = 0.15) [4]. Furthermore, Dahl and Akerud disclosed that the on-target side effect could be reduced by reducing the drug residence time [5]. Thus, a drug with optimal efficacy and toxicity profile should have a balanced k_on_ and k_off_. Since IC_50_ and K_d_ depend on the measurement of the combined effect of k_on_ and k_off_, they are actually insufficient to explain the impact of binding/unbinding kinetic on drug action, as the same value of K_d_ can come from infinite number of combinations of k_on_ and k_off_. Additionally, since K_d_ is dependent on the free energy difference between the bound and unbound states but is independent on the transition state of protein-ligand interaction (PLI), it is inadequate to explain the binding/unbinding process of PLI [4,6].

Experimental techniques for the study of PLI kinetics are not only expensive and time-consuming but also insufficient of providing detailed molecular characterization of the PLI kinetics process [7–9]. Computational modeling plays an increasing role in elucidating the binding/unbinding process of PLI. Molecular dynamics (MD) simulations have been reported to be capable to capture the binding process, from beginning to end, in full atomic detail [10]. Unfortunately, the power of MD simulations is limited due to the fact that protein-ligand binding event takes place in a time scale ranging from microseconds up to hours and days. For the majority of the binding processes, they are infeasible for MD simulations. Therefore, metadynamics and other conformational sampling techniques have been developed to improve sampling in MD simulations of a system where ergodicity is hindered by the form of the system’s energy landscape. Gervasio et al. applied a metadynamics method successfully to the docking of ligands on flexible receptors in water solution. The method is able not only to find the docked geometry and to predict the binding affinity (ΔG_binding_) but also to explore the entire docking process from the solution to the docking cavity, including barriers and intermediate minima [11]. Even though these progresses are remarkable, metadynamics is not yet feasible to study the whole binding/unbinding process of PLI on a large scale. In addition, the choice of collative variables in the metadynamics simulation is not a trivial task.

With the increasing availability of protein binding kinetics data [12,13], data-driven modeling provides an alternative and efficient solution to studying the PLI kinetics. Several predictive models for kinetic constants of protein-protein interaction (PPI) have been developed [14,15]. However, the molecular attributes in these models only covered static structural characteristics such as the percentage of residues in α-helix, the buried surface area of protein, the proportion of charged residues and the proportion of polar atoms at the interface, and the energetic features such as hydrogen bonding potential and the interfacial electrostatic interaction energy between interfacial residues. These features may not sufficiently capture conformational dynamics of the PLI kinetic processes. In addition, existing methods predict k_on_ and k_off_ independently. As a matter of fact, they could be dependent in nature. To our knowledge, few methods are available for the large-scale modeling of the binding/unbinding kinetics of PLI with explicit dynamic features, as well as predicting k_on_ and k_off_ simultaneously.

To tackle the above problems, we integrate energetic and conformational dynamic features derived from efficient molecular modeling with state-of-the-art multi-task learning (MTL) approach. In this study, ligand-bound HIV-1 proteases are used as an example to build models. In addition to Electrostatic Energy (EE) and van der Waals Energy (VDWE), which are derived from all-atom Molecular Dynamics simulation [16,17] and environmental-dependent electrostatic potential energy [18], Relative Movement of Ligand-Residue (RMLR) and Relative Movement of Residue-Residue (RMRR) that represent the dynamics impact of ligand binding on the amino acid residues are derived from Normal Mode Analysis (NMA) analysis and used to train machine learning models. Our multi-facet statistical analysis consistently shows that conformational dynamic features, such as RMLR, are as important as energetic features, particularly EE, in predicting k_on_ and k_off_. Based on these findings, we propose that coherent conformational dynamic coupling between protein and ligand may play a critical role in determining the kinetic rate constants of PLI. Furthermore, we demonstrated that NMA is an efficient method to capture conformational dynamic features of the binding/unbinding kinetics of PLI. Coupled with the state-of-the-art multi-target classification as well as multi-target regression, it is possible for us to screen and optimize compounds based on the binding/unbinding kinetics of PLI in a high-throughput fashion. The further development of such computational tools will bridge one of the critical missing links between *in vitro* drug potency and *in vivo* drug efficacy and safety, thereby accelerating drug discovery process.

## Results

### 1. Characteristics of data set

In this study, we used HIV-1 protease complex structure to investigate the conformational dynamics and develop predictive model of ligand binding/unbinding kinetics. The HIV-1 protease is an excellent model system for our purpose. First, thirty-nine HIV-1 protease inhibitors have experimentally determined k_on_ and k_off_ under the same condition [19,20]. They provide reasonable number of high quality data points for the data-driven modeling. Second, abundant data of HIV-1 protease inhibitor resistance mutation (PIRM) are available [21]. They can be used to validate the predictive model. Third, both unbound and complex structures of HIV-1 protease are released in Protein Data Bank [22]. The apo- and holo-conformations are the basis for our analysis.

When mapping the 39 HIV-1 protease inhibitors on the 2-dimensional space of k_on_ and k_off_, as shown in Figure 1, all FDA-approved drugs were clustered in the upper-left corner with high k_on_ and low k_off_. Based on the criteria of log_10_k_off_ = -2 and log_10_k_on_ = 5.6, which will put all FDA approved drugs in a single class and evenly distribute the inhibitors into four different classes, with the labels (0,0), (0,1), (1,0), and (1,1) (see Supplementary Table S1). It is noted that several inhibitors such as A037 have the similar value of K_d_, which is equal to k_off_/k_on_, to that of the approved drugs, but fall into different classes from the FDA-approved drugs in the 2D map. It suggests that atomic interactive constant Kd alone is not sufficient to determine the drug effect.

**Figure 1.**
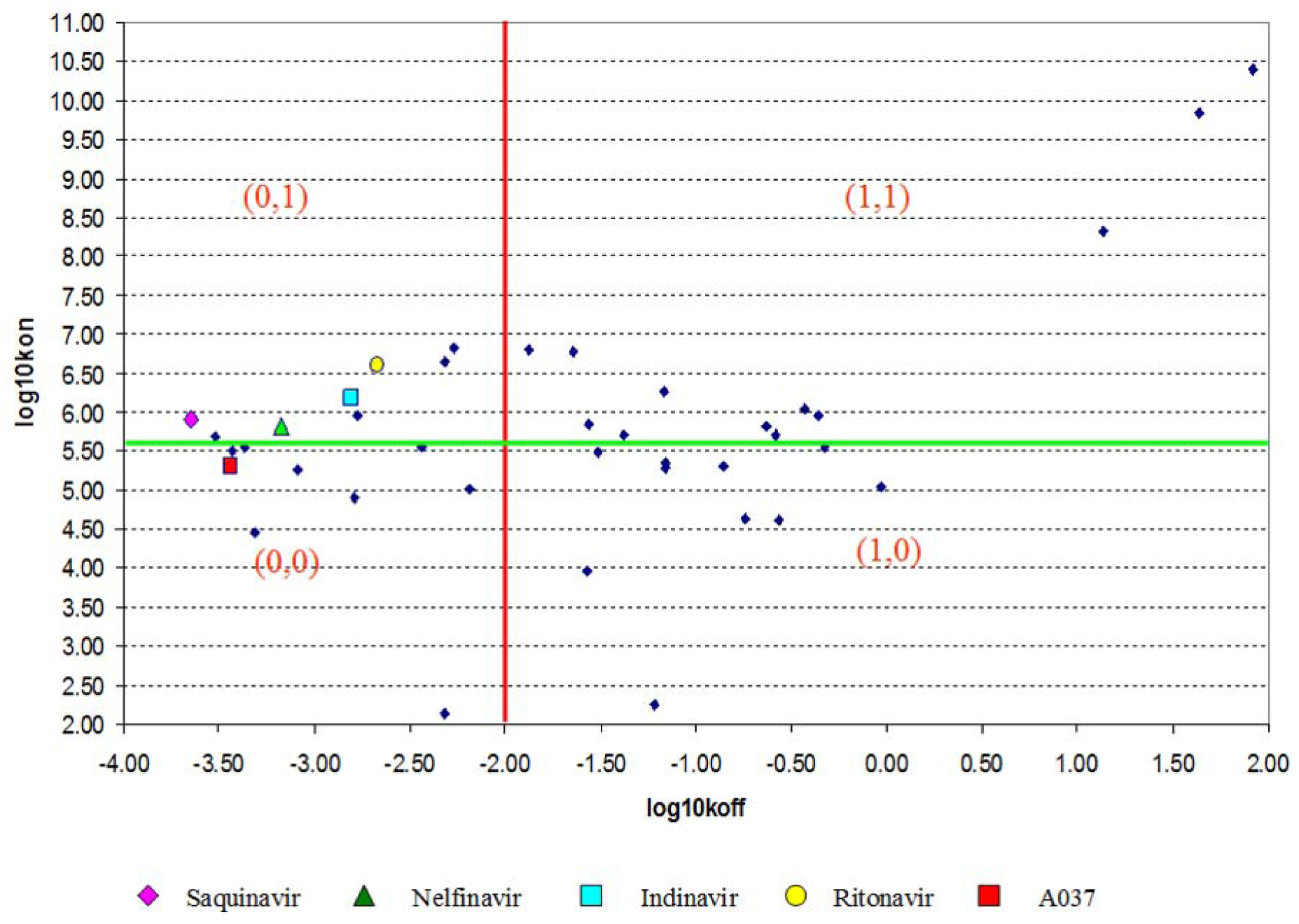
Discretization of k_on_ and k_off_ of HIV protease inhibitors. Results of the discretization based on the criteria set at log_10_k_off_ = -2 (x-axis) and log_10_k_on_ = 5.6 (y-axis). Thirty-nine training records were discretized into four binary classes: (0,0), (0,1), (1,0), and (1,1).

Ten inhibitors have solved HIV-1 complex structures in PDB. For the remaining inhibitors whose complex structures have not been experimentally determined, protein-ligand docking software eHiTS [23] is applied to predict its binding pose. The receptor is chosen from one of the ligand-bound HIV-1 complexes with the co-crystallized ligand structure similar to the docked ligand structure. Whenever possible, the common fragment of the co-crystallized and the docked ligand is used as a constraint to select the final binding pose of the docked ligand, such that the RMSD of superimposed common fragments is minimal. An example is shown in Supplementary Figure S1.

Binding site amino acid residues that are involved in the HIV-1 protease inhibitor interactions are determined using the change of solvent assessable surface area (SASA) upon ligand binding. As depicted on Figure S2, there are total 44 amino acids on both chains of the HIV-1 dimmer.

### 2. Characterization of protein-ligand interaction using the directionality of normal modes

Normal Mode Analysis (NMA) is a powerful computational method to identify and characterize the slowest molecular deformational motions with large amplitude, which are widely involved in biological functions of macromolecules, but inaccessible by other methods. Protein binding and unbinding events are often on a long-time scale ranging from milliseconds to days, far beyond the current capability of MD simulations. Coarse-grained NMA may allow us to extract important dynamic information on protein-ligand binding/unbinding processes. Since the presence of solvent damping dramatically slows down the large-amplitude motions of biomolecules, the timescales of molecular motions in reality are much longer than what can be estimated from the eigenvalues of NMA that are calculated in vacuum. In other words, solvent damping causes a discrepancy on a timescale between NMA and real molecular motions. However, the study conducted by Ma revealed that the presence of solvent has a minor impact on eigenvectors, which are determined by the potential surface only [24]. Thus, the information provided by the eigenvectors for the directionality of conformational transitions could be used to study dynamic processes in the time-scale of real situations.

In this study, NMA was conducted using iMod [25]. The directionality of normal modes of the residues in the binding site is used to characterize the conformational dynamic features of binding and unbinding event. Specifically, two data sets including Relative Movement of Ligand-Residue (DS-RMLR), and Relative Movement of Residue-Residue (DS-RMRR) were derived from NMA analysis. Both DS-RMLR and DS-RMRR cover the 10 lowest frequency modes, where DS-RMLR illustrates the relative directionality of normal modes between ligand and residue, and DS-RMRR illustrates the change of directionality of normal modes of binding site residues upon the ligand binding. As an example, Figure 2A depicts the superposition of the 44 residue eigenvectors of the aligned apo structure and the DMP bound structure of 1^st^ normal mode. It illustrates the shift of the eigenvectors of the 44 residues of HIV-1 protease upon ligand binding. Figure 2B illustrates the relative displacements of the 44 ligand-residue pairs in the DMP bound HIV-1 complex.

**Figure 2.**
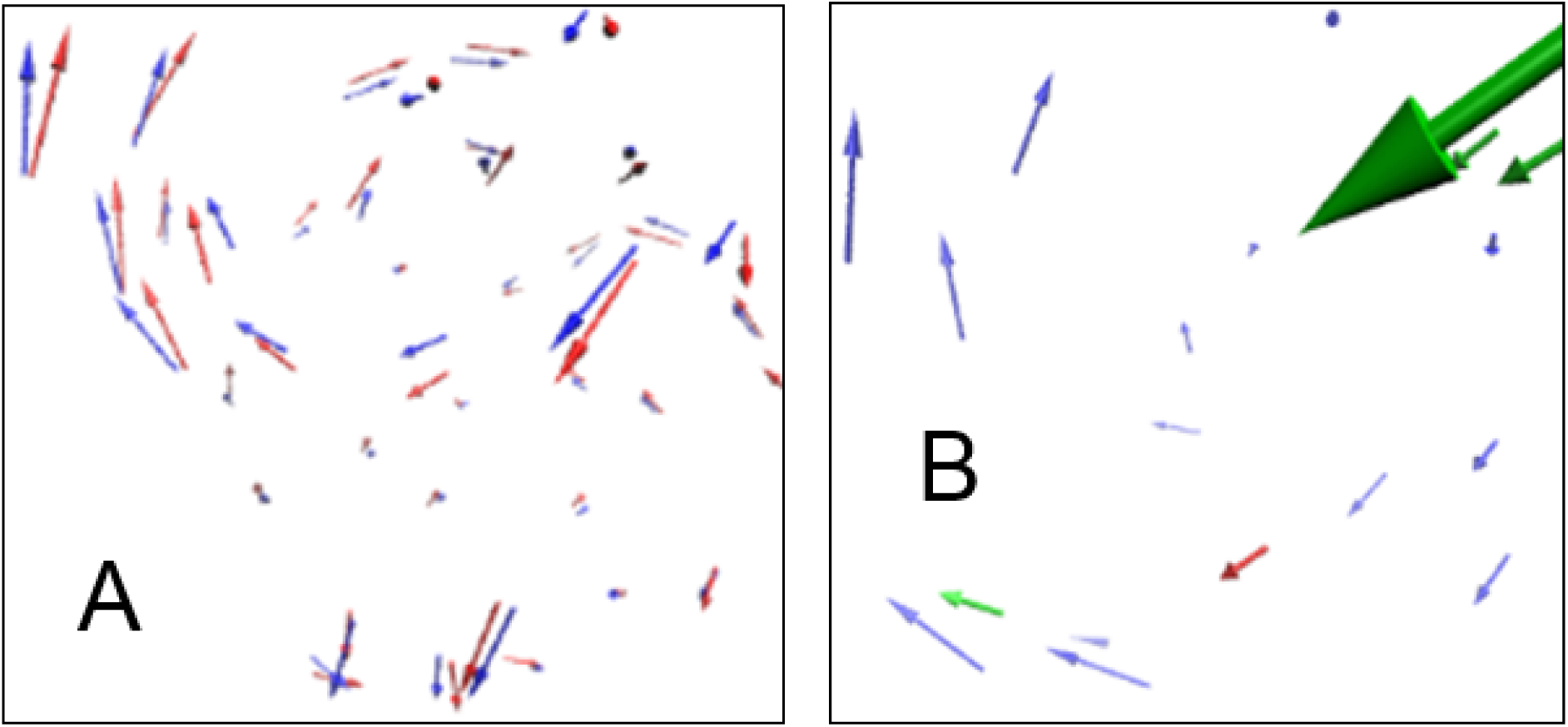
Directionality of normal mode. (A) Superposition of the 44 residue eigenvectors of the aligned apo HIV-1 structure (red) (PDB code:3IXO) and the DMP bound HIV-1 structure (blue) (PDB code: 1QBS) of 1^st^ normal mode. (B) Eigenvector displacements of the 44 DMP (red) –residue (green/blue) pairs in the DMP bound HIV-1 complex (1^st^ normal mode). Green/blue arrows are the eigenvectors of the 22 residues of chain A/B respectively.

### 3. Characterization of ligand-residue interaction energy

Residue decomposed Pairwise Interaction Energy (PIE) and its two constituting components including Electrostatic Energy (EE) and van der Waals Energy (VDWE), between the ligand and the binding site residue of HIV-1 protease, are calculated from all-atom Molecular Dynamics (MD) simulation and environmental-dependent electrostatic potential energy. The values of PIE, EE, and VDWE, which characterize various energetic aspects of ligand-residue interaction, are used to build three data sets: DS-PIE, DS-EE, and DS-VDWE.

### 4. Structural determinants of protein-ligand binding/unbinding

We use the energetic and conformational dynamic attributes derived from MD simulation and NMA to train a multi-target machine learning (MTML) model for the classification prediction of kinetic rate constants. In total, there are five principal training data sets including DS-PIE, DS-EE, DS-VDWE, DS-RMLR, and DS-RMRR. Each of them comprises thirty-nine cases with each case comprising 44 attributes.

MTML is defined as follows: Given a set of learning examples D of the form (X,Y), where X = (x_1_, x_2_,…, x_k_) is a vector of k training attributes and Y = (y_1_, y_2_,…, y_t_) is a vector of *t* target attributes, learn a model that, given a new unlabeled example X, can predict the values of all target attributes Y simultaneously. When y_i_ is categorical, the problem is known as classification. In this study, the yi is a binarized value of k_on_ and k_off_ as shown in Figure 1.

Random Forest Predictive Clustering (RF-Clus) is applied for the task of MTML. RF-Clus outperforms other MTML algorithms in the benchmark studies [26]. In addition, it can handle high-dimensional features, e.g. in the situation where the number of attributes is much higher than the number of cases, and can select the importance of attributes (amino acid residues) that contribute to the accuracy of k_on_/k_off_ prediction. The model was run on the iteration numbers of 100, 200, 250, and 500 in the leave-one-out cross-validation experiment.

Table 1 shows the selected features in the descending order of score of importance. Consequently, sixteen, fifteen, thirteen, and fourteen features were selected from DS-RMLR, DS-RMRR, and DS-EE, and DS-PIE. These identified key residues consist of three motifs: an N-terminal motif (R8, L10), a charged motif (L23, D25, G27, A28, D29, D30, and V32), and a motif corresponding to flap region (residue 43-58), as shown in Figure 3. Both the N-terminal motif and the charged motif are common to DS-PIE, DS-RMLR, DS-RMRR and DS-EE. The flap region is identified by DS-RMRL and DS-RMRR.

**Table 1.** Key residues identified from four data sets: DS-RMLR, DS-RMRR, DS-EE, and DS-PIE.

| Residue | DS-RMLR |  | DS-RMRR |  | DS-EE |  | DS-PIE |  |
| --- | --- | --- | --- | --- | --- | --- | --- | --- |
|  | Frequency | Score | Frequency | Score | Frequency | Score | Frequency | Score |
| R8 | 35.86 | 0.78 | 56.46 | 0.73 | 64.79 | 0.74 | 52.32 | 0.73 |
| L10<sup>†</sup> | 39.13 | 0.75 | 61.16 | 0.71 | 34.72 | 0.77 | 43.04 | 0.75 |
| L23<sup>†</sup> | 45.60 | 0.71 | 54.18 | 0.71 | 40.69 | 0.72 | 35.52 | 0.76 |
| D25 | 44.21 | 0.69 | 45.02 | 0.73 | 37.41 | 0.75 | 49.28 | 0.70 |
| G27 | 42.85 | 0.68 | 39.96 | 0.73 | 30.62 | 0.75 | 35.40 | 0.72 |
| A28 | 34.70 | 0.68 | 39.67 | 0.70 | 35.23 | 0.71 | 27.48 | 0.75 |
| D29 | 28.18 | 0.71 | 41.37 | 0.67 | 28.36 | 0.73 | 48.60 | 0.67 |
| D30<sup>†</sup> | 30.95 | 0.70 | 39.79 | 0.67 | 25.90 | 0.72 | 28.24 | 0.71 |
| V32<sup>†</sup> | 29.35 | 0.66 | 37.83 | 0.66 | 25.15 | 0.72 | 29.84 | 0.70 |
| K45 | 37.77 | 0.65 | 35.79 | 0.68 |  |  | 27.98 | 0.70 |
| I47<sup>†</sup> | 30.44 | 0.68 | 35.19 | 0.66 | 25.54 | 0.68 |  |  |
| G48<sup>†</sup> | 26.88 | 0.67 | 26.10 | 0.66 |  |  |  |  |
| G49 | 34.44 | 0.63 | 31.57 | 0.63 | 33.23 | 0.66 |  |  |
| I50<sup>†</sup> |  |  | 36.64 | 0.61 |  |  |  |  |
| A52 | 26.08 | 0.66 | 27.07 | 0.62 |  |  | 26.73 | 0.64 |
| F53<sup>†</sup> | 27.90 | 0.66 |  |  |  |  |  |  |
| L76<sup>†</sup> |  |  |  |  |  |  | 26.63 | 0.62 |
| P81 |  |  |  |  | 25.54 | 0.63 | 25.78 | 0.61 |
| R8* | 30.55 | 0.61 |  |  |  |  |  |  |
| D25* |  |  |  |  |  |  | 31.25 | 0.60 |
| D29* |  |  |  |  | 28.13 | 0.65 |  |  |

The feature selection criterion is the frequency of attribute occurrence $\geq 25\%$ .
\*Residues in chain B.
<sup>†</sup>Residues whose mutation lead to drug resistance.

**Figure 3.**
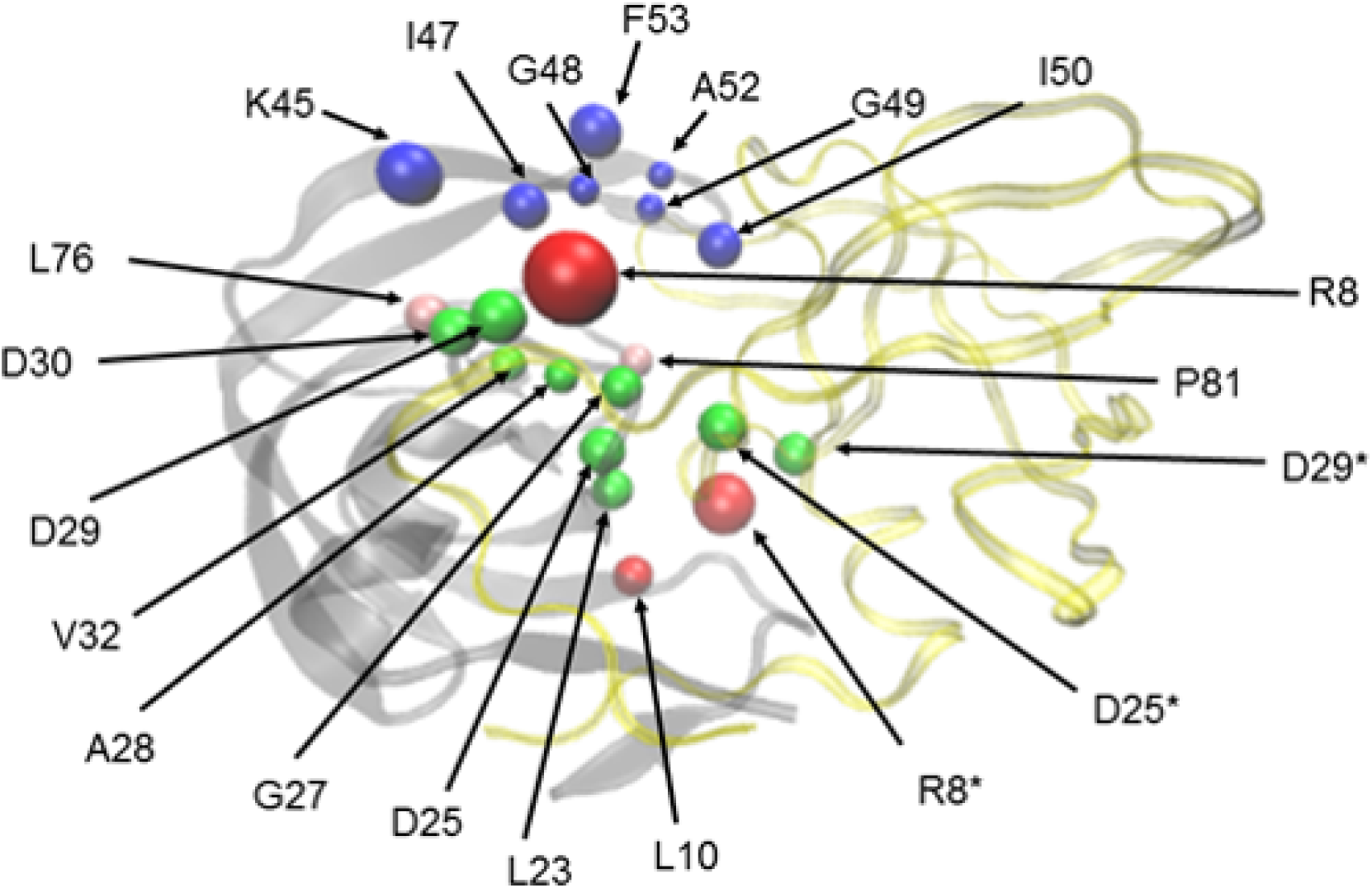
Twenty-one key residues. Chains A and B of HIV-1 protease are in transparent grey and transparent yellow ribbons, respectively. The three residues located on the chain B are labeled with * superscript. Charged motif including L23, D25, G27, A28, D29, D30 and V32 are in green. Residues located on the N-terminal including R8 and L10 are in red. Residues located in the flap region including K45, I47, G48, G49, I50, A52, and F53 are in blue; L76 and P81 located near the flap region are in pink.

All-atom MD simulations have shown that the conformational dynamics of flap region (residue 43-58) plays a key role in the ligand binding process of HIV-1 protease [27,28,29]. Consistent with this observation, the residues in the flap region are identified as key kinetic features with signification displacement upon ligand binding. The recapitulation of the findings from the MD simulation provides a validation to the data-driven approach in the paper. The charged motif in the active site participates in substrate peptide recognition. Specifically, D25 and D29 form hydrogen bonds with substrate peptide. Additionally, R8 and D30 can interact with polar side chains or distal main chain groups in longer substrate peptides. Moreover, the mutation of L10, L23, and V32 lead to drug resistance of HIV protease inhibitors.

### 5. Combined electrostatic and conformational dynamic features can predict k_on_/k_off_ accurately

We examine the impacts of the energetic and conformational dynamic features on the prediction of k_on_/k_off_ accuracy by building different MTML models in three stages. First, we use energetic features (data sets: DS-EE, DS-PIE, DS-VDWE) to build the MTML model. Second, in order to evaluate if the normal mode directionality features can be used to predict the ligand binding and unbinding process, we apply DS-RMRR, DS-RMLR and DS-RMLR+DS-RMRR comprising 88 training attributes in the feature vectors to build the MTML model. Third, we integrate the properties of conformational dynamics and energetics by adding the RMLR features to DS-EE (data set DS-EE+DS-RMLR) to build the MTML model.

For the models trained by DS-EE, DS-PIE, DS-VDWE, DS-RMLR, DS-RMRR, DS-RMLR+DS-RMRR, and DS-EE+DS-RMLR, the highest prediction accuracy of log_10_k_on_ are 71.79, 69.23, 43.59, 69.23, 51.28, 69.23, and 76.92% respectively (Figure 4A), the highest prediction accuracy of log_10_k_off_ are 76.92, 66.67, 56.41, 71.79, 64.10, 71.79, and 71.79% respectively (Figure 4B), and the highest prediction accuracy of the combined four-class log_10_k_on_/log_10_k_off_ are 71.79, 66.66, 47,43, 69.23, 57.69, 70.51, and 74.35% respectively (Figure 4C).

**Figure 4.**
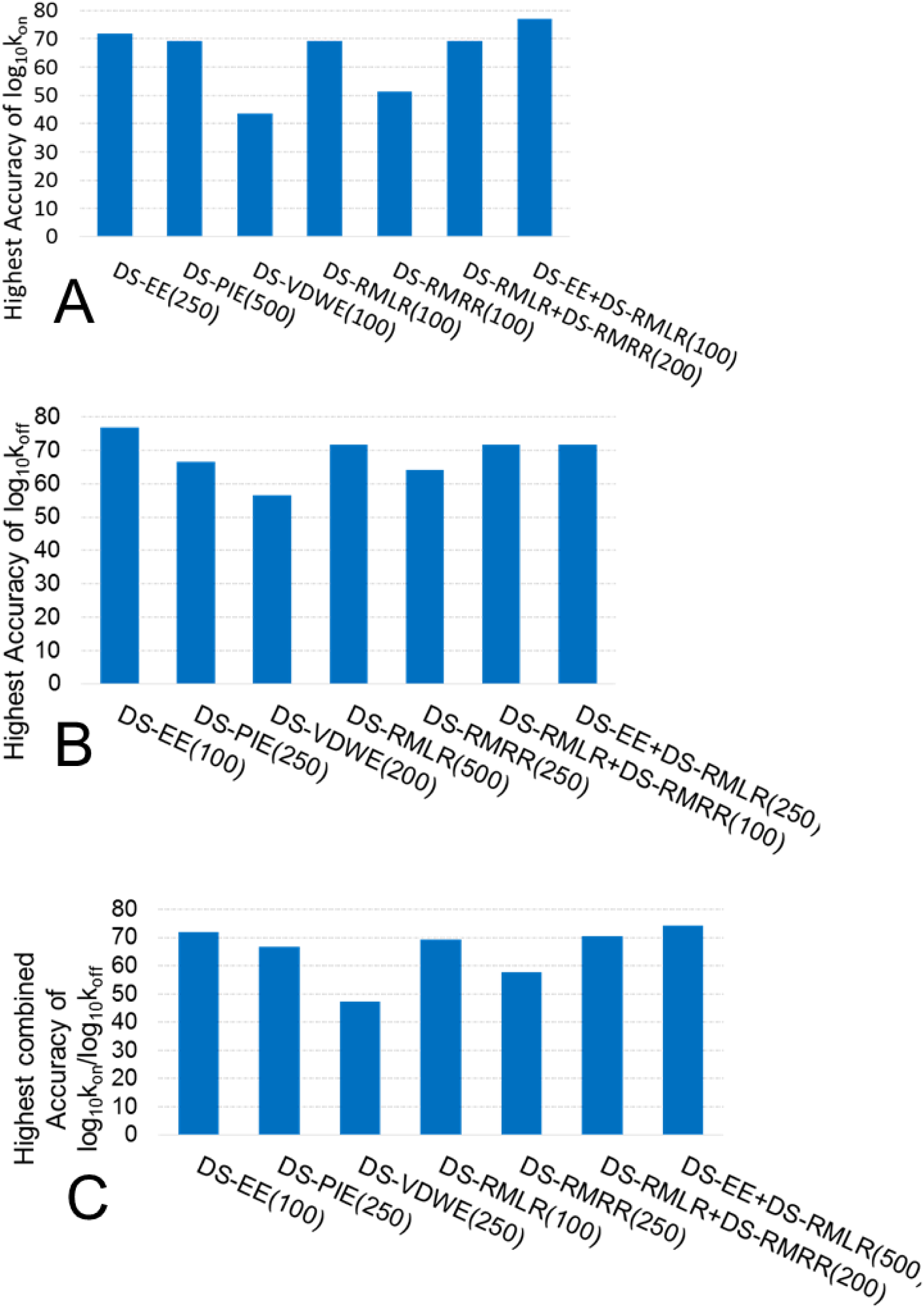
Prediction accuracy of MTML model. Prediction accuracy of (A) log_10_k_on_, (B) log_10_k_off_, and (C) the combined four-class log_10_k_on_/log_10_k_off_. The number in parentheses is the iteration number used in the experiment.

Among the three models trained by the energetic features, the prediction accuracy of the combined four-class log_10_k_on_/log_10_k_off_· given by the DS-EE and DS-PIE models are significantly higher than a random guess (50%) by 21.79 and 16.66% respectively, but the accuracy given by the DS-VDWE model is lower than random by 2.57%. These results suggest that in the case of HIV-1 protease, electrostatic interaction plays a key role in the binding/unbinding process, and the Electrostatic Energy features are more accurate in predicting k_on_ and k_off_ than the features of van der Waals Energy and Pairwise Interaction Energy.

For all the three models trained by the normal mode directionality features, the prediction accuracy of the combined four-class log_10_k_on_/log_10_k_off_ is higher than random. Although the accuracy given by the DS-RMRR model is only slightly higher than random by 7.69%, the accuracy given by the DS-RMLR and DS-RMLR+DS-RMRR models are significantly higher than random by 19.23, and 10.51% respectively. These results suggest that the normal mode directionality can capture the information on the ligand binding and unbinding process.

Comparing with the prediction accuracy of the combined four-class log_10_k_on_/log_10_k_off_ given by the DS-EE and DS-EE+DS-RMLR models shows that integrating the conformational dynamic features into the energetic features increases the accuracy from 71.79 to 74.35 by 2.56%. Consequently, it implies that the electrostatic interaction and conformational dynamics are jointly responsible for the binding kinetics of HIV protease.

## Discussions

### 1. Coherent receptor-ligand movement is one of the structural determinants of protein binding/unbinding kinetics

Consistent with the all-atom MD simulation, the MTML model trained with the relative directionality of normal mode between residue and ligand recapitulates the role of flap region in the binding kinetics of HIV protease. It is known that electrostatic interaction between a charged drug and a charged receptor impacts the kinetic rate constants [30,31]. Specifically, k_on_ is sensitive to long-range electrostatic interaction, and k_off_ tends to be influenced more by shortrange interactions such as hydrogen bonds, salt bridges and van der Waals contacts [32]. The majority of the residues selected in this study are hydrophobic; the exceptions are the catalytic D25 and D29, which are able to form hydrogen bonds with the main chain groups of substrate peptides, and R8, D30 and K45 which can interact with polar side chains or distal main groups in longer substrate peptides. The MTML model ranks these charged residues more important than the flap region in their contribution to the prediction accuracy. In addition, the MTML model can achieve high prediction accuracy using the electrostatic energy alone. These indicate that the electrostatic interaction is one of the major factors in determining the binding/unbinding kinetics of HIV protease.

Interestingly, in addition to the electrostatic interactions, the directionality of ligand binding site residue movement also has strong correlations with the kinetic constants. Not only the similar residues are selected from DS-RMLR to those from DS-EE, the best performed MTML model is obtained from the combination of DS-RMLR and DS-EE. Based on this observation, we propose that the coherent movement between the ligand and the receptor may play a critical role in determining the ligand binding and unbinding kinetics. As shown in Figure 5, even if two protein-ligand complexes have the same non-covalent interactions with the same intensity, they may have different kinetic constants due to the different relative movements between the ligand atom and the receptor atom. It is not surprising, as the non-covalent interactions, especially hydrogen-bonding, depends on the relative directionality of atomic pairs. The relative movement may change the directionality of the interaction, thus weaken (even break) or strengthen the interaction. Thus, the coherent conformational dynamics coupling could be one of key structural determinants of protein binding/unbinding kinetics. This has not been observed before.

**Figure 5.**
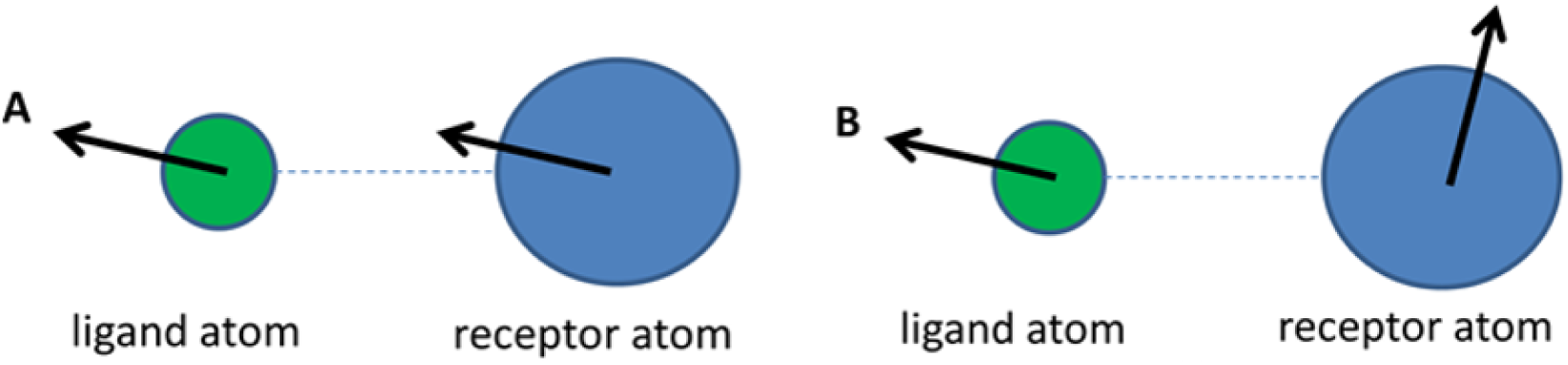
The influence of ligand-residue conformational coupling on protein binding/unbinding kinetics. (A) Coherent conformational coupling. The relative movement between ligand atom and receptor atom will not change the distance and directionality of the interaction, thus the intensity of interaction will not be changed. (B) Incoherent conformational coupling. The relative movement between ligand atom and receptor atom will alter the distance and directionality of the interaction. As a result, the interaction could be weaken or broken.

### 2. High-throughput predictive modeling of ligand binding and unbinding kinetics

In spite of recognized importance of protein-ligand binding and unbinding kinetics in the drug discovery, few efficient computational tools are available to screen and optimize chemical compounds based on the binding and unbinding kinetics. With the increasing availability of experimentally determined k_on_/k_off_ data [14,15], data-driven approach is an appropriate choice for the development of a high-throughput predictive model of ligand binding and unbinding kinetics [33]. However, two questions remain to be answered in developing an effective and efficient machine learning model. First, what are the molecular determinants of ligand binding and unbinding kinetics so that they can be used as features to train a high-quality machine learning model with the minimum impact of over-fitting, and false correlation? Second, what are the suitable machine learning algorithms that can handle high-dimensional data and predict k_on_/k_off_ simultaneously? For the first time, we have shown that NMA could be an efficient tool to capture the conformationally dynamic information of the ligand binding and unbinding kinetics. The features derived from the NMA could be used to enhance the performance of the machine learning model. Moreover, recently developed multi-target classification algorithms such as RF-Clus could be adopted to train a machine learning model that can predict dependent k_on_/k_off_ simultaneously.

Although this proof-of-concept study demonstrates the potential of integrating physically-based modeling with multi-target machine learning in understanding the molecular determinants, and developing high-throughput predictive model of ligand binding and unbinding kinetics, there is plenty of space to improve the methodology. Since solvation effect causes a discrepancy on a timescale between real molecular motion and NMA that are calculated in vacuum, it is expected that NMA coupled with an implicit or explicit solvation model may provide more information on the conformational dynamics of ligand binding process. As water plays a critical role in the ligand binding, the explicit incorporation of the water molecule in the binding site may improve the accuracy of simulation. The global and local geometry of binding pocket could be another important feature [34,35]. In the current study, the ligand is treated as a single rigid body. As a matter of fact, the flexibility of the ligands may have impacts on the kinetic rate constants. As shown in supplemental information Figure S3, both of the values of log_10_k_on_ and log_10_k_off_ are weakly correlated with the ligand flexibility that is characterized by the number of rotatable bonds. The general trend is that the k_on_ and k_off_ decrease as the number of the ligand rotatable bonds increases. It suggests that the performance of MTML model could be further improved by incorporating the ligand properties. We group the k_on_/k_off_ into four classes and use the classification model to predict the class and to select features. In practice, it could be more useful to predict the real value of kon and koff simultaneously. It requires a multi-target regression model, which is an active area of research in machine learning.

There are three different models of conformational ensemble of protein-ligand complex. They are the model of induced fit mechanism which is adopted by HIV-1 protein-ligand complex [1,36], the model of selected fit mechanism [37,38], and the model of three step mechanism [39,40]. The mechanism of model determines the on-rate and off-rate equations. For example, the induced fit on-rate is limited by the diffusional rate of encounter complex formation of the proteins in their unbound conformational ensemble, but the off-rate is dependent on the equilibrium between the ground state complex and the excited state complex [36]. Since all the training data sets in this study only cover the characteristics of the ground state HIV-1 complex, the ignorance of the characteristics of the excited state HIV-1 complex could induce deficiency in the predictive model. In summary, the further development of predictive modeling tools of ligand binding and unbinding kinetics will bridge one of the critical missing links between in vitro drug potency and in vivo drug efficacy and safety on a large scale, thereby accelerating drug discovery process.

## Materials and Methods

Figure S4 depicts the workflow of computational procedure in this study, which includes four phases: Phase 1 concerns the structure construction of 3D ligand-bound HIV-1 protease complex. Phase 2 addresses the identification of ligand binding site residues. Phase 3 targets the construction of the five principal data sets. Phase 4 is machine learning computation.

In brief, the 3D conformation of ligand was docked in the HIV protease if no cocrystallized structures exist. Normal Mode Analysis (NMA) was performed for both apo- and holo-structure for each inhibitor. Relative Movement of Ligand-Residue (RMLR) and Relative Movement of Residue-Residue (RMRR) that represent the conformational dynamics impact of ligand binding on the binding site residues were derived from NMA analysis. In addition, Pairwise Interaction Energy as well as its two components, van der Waals Energy and Electrostatic Energy between the ligand and amino acid residues, were derived from the 200 ps all-atom Molecular Dynamics simulation and environmental-dependent electrostatic potential energy. Finally, conformational dynamics and thermodynamics features, individually or combined, are used to train multi-target machine learning models.

## Supporting Information^*^

### SI Materials and Methods

In this study, thirty-nine ligand-bound HIV-1 protease complexes were used as training samples to build five principal datasets including three datasets with energetic features: Electrostatic Energy (DS-EE), van der Waals Energy (DS-VDWE), and Pairwise Interaction energy (DS-PIE) that is the sum of EE and VDWE, and two datasets with conformational dynamic features: Relative Movement of Ligand and Residue (DS-RMLR), and Relative Movement of Residue-Residue (DS-RMRR). Each of them comprises 39 cases with each case comprising 44 attributes. Two datasets with 88 training attributes including DS-RMLR+DS-RMRR and DS-EE+DS-RMLR were built by integrating DS-RMLR and DS-RMRR, and DS-EE and DS-RMLR respectively. Thus, a total of seven datasets was used to train the RF-Clus MTML model for classification prediction to predict k_on_ and k_off_ simultaneously.

Figure S4 depicts the workflow of computational procedure in this study, which includes four phases: Phase 1 concerns the structure construction of 3D ligand-bound HIV-1 protease complex. Phase 2 addresses the identification of residues that are close to the ligand. Phase 3 targets the construction of the three principal datasets. Phase 4 is machine learning computation.

### Phase 1 - 3D Structure of Ligand-Bound HIV-1 Complex

In 2002, Markgren et al. reported the kinetic rate constants (k_on_ and k_off_) of thirty-nine ligand-bound HIV-1 complexes [19] using the technique of Surface Plasmon Resonance Based (SPR) Biosensor (Table S2) [20]. Thirty-three of them were classified into five structural categories (Table S3) in reference to the 2D molecular structure of B206 as shown in Figure S5. The five categories include non-B268 analogues, P1/P1’ analogues of B268, P2/P2’ analogues of B268, cyclic ureas and cyclic sulfamides. Standard nomenclature, P_1_…P_n_, P_1_’…P_n_’ is used to designate amino acid residues of peptide substrates in the enzyme-substrate interactions (Figure S6). In this study, ten 3D molecular structures of the thirty-nine complexes were collected from RCSB Protein Data Bank (PDB) [41]. They include DMP-1QBS, AMP-3EKV, B435–1D4H, B369-1EBY, B409-1EC1, B388-1EBZ, B425-1D4I, Nelfinavir-3EKX, Ritonavir-1HXW, and U75875-1HIV. The remaining twenty-nine complex structures were obtained from a three-step building process. First, a 2D molecular ligand structure was transformed into its SMILES string [42]. Figure S7 depicts the transformation from the 2D molecular structure of B268 into its SMILES string. Second, the ligand SMILES string was converted into a 3D molecular ligand structure using Frog [43] as shown in Figure S8. Third, Electronic High Throughput Screening (eHiTS) program [23,44-46] was used to dock a target ligand into the active site of a wild type HIV-1 protein. The receptor was chosen from one of the five ligand-bound HIV-1 complexes (1QBS, 1EBW, 1AJV, 1EC2, and 1D4H) with the co-crystallized ligand structure similar to the target ligand structure. Whenever possible, the common fragment of the co-crystallized and the docked ligand is used as a constraint to select the final binding pose of the docked ligand, such that the RMSD of superimposed common fragments is minimal. An example is shown in Figure S1. Table S4 shows the results of the docking process.

**Detection Limits of SPR Biosensor:** Due to the baseline stability of SPR biosensor and the detection limit exerted by the diffusion rate of an ligand to its binding partner which is immobilized on the biosensor surface, SPR biosensor is only capable of measuring association (k_on_) and dissociation (k_off_) kinetic rate constants in the range of 10^2^ to 10^8^ M^−1^s^−1^ and 1 to 10^−6^s^−1^, respectively [7]. As shown in Table S2, the k_on_ and k_off_ values of DMP, B376, and A008 are beyond the upper detection limit. The k_on_ values of B277 and A016 are proximate to the lower detection limit.

### Phase 2 Identify Residues close to Ligand

Solvent accessible surface area (SASA) procedures [47] with probe radii of 1.4 and 2.1Å, written in TCL script were implemented on Visual Molecular Dynamics (VMD) [48] platform to identify residues close to the ligand in a ligand-protein complex. The steps of the SASA computation are as follows: First, calculate the SASA for each residue in the ligand-protein complex. Second, calculate the SASA for each residue in the isolated protein molecule. Third, subtract the results from the above two steps for the same residue. Only residues that are close to the ligand have non-zero values after subtraction. The results reveal that 44 residues out of 198 are close to the ligands within 4.2 Å. Figure S9 shows the TCL script of SASA with probe radius of 1.4Ä for the A045-1AJV complex.

### Phase 3 Principal Data set Construction

Five principal training data sets including DS-PIE, DS-VDWE, DS-EE, DS-RMLR, and DS-RMRR were constructed for the ML prediction of kinetic rate constants (log_10_k_on_ and log_10_k_off_). Normal mode analysis was performed to compute the attribute values in DS-RMLR and DS-RMRR.

1. **Training data set DS-PIE:** DS-PIE covers the PIE properties of a residue-ligand pair. The energy comprises two terms: van der Waals Energy + Electrostatic Energy (kcal/mol)

- van der Waals Energy = C_12_/r^12^ – C_6_/r^6^, where r is the distance between the two atoms’ nuclei, C_12_ and C_6_ are constants, whose values depend on the depth of the energy well and the equilibrium separation of the two atoms' nuclei.
- Electrostatic Energy = Cq_i_q_j_/εr_ij_, where C is the Coulomb constant; q_i_ and q_j_ are point charges i and j; ε is the dielectric constant; and r_ij_ is the distance between the two points. Since ε can range from 1 to 80 in a protein environment, a reasonable ε value is important to the correctness of the electrostatic energy calculation, and determines the accuracy of the PIE calculation. Average dielectric constants of different types of residues (Figure S10) reported by Li et al. [18], were adopted in this study to calculate the electrostatic energy. The dielectric constants range from 11.0 to 25.6 and are physically sound. Charged amino acids (Lys, Arg, Glu, and Asp) are associated with the highest average dielectric values. They also tend to be loosely packed on the protein surface, leaving room for structural rearrangement. On the other hand, hydrophobic residues (Cys, Ile, Phe, Val) are assigned with low dielectric values and they tend to be found in the protein core.

MD simulations of the thirty-nine ligand-HIV-1 models were carried out using the Nanoscale Molecular Dynamics (NAMD) [17] program with CHARMM27 force field for HIV-1 protein and CHARMM general force field for the ligands [49]. All models proceeded through a minimization process of at least 4 ps and an equilibrium process of 200 ps. During the simulations, temperature was set at 310 K, and the generalized born implicit solvent method was used with the ionConcentration, GBISDelta, GBISBeta, GBISGamma and alphaCutoff set at 0.15, 0.8, 0.0, 2.90912 and 14, respectively [50-52]. After loading the 200 ps trajectory file produced from MD simulations into VMD, the value of PIE of each unique residue-ligand pair was calculated using the NAMDEnergy plugin in VMD. Figure S11 shows the NAMDEnergy graphical user interface for the computation of the PIE between ligand A045 and leucine of chain A. The expression of “(chain A) and (resid 10)” on the tab of Selection 1 identifies leucine of chain A with the dielectric constant of 11.8 entered on the dielectric tab, and the expression of “resid 501” on the tab of Selection 2 identifies ligand A045.

MD simulations were performed at the High Performance Computing Center at the College of Staten Island, CUNY.

2. **Data sets DS-RMLS and DS-RMSS:** The training attributes of DS-RMLR and DS-RMRR illustrate the relative movement of a ligand-residue pair and the relative movement of a residue upon ligand binding respectively. The NMA system, iMOD [25] developed by Lopez-Blanco et al. in 2011 was adopted to compute the displacement vectors of the residues and the ligands in the models. iMOD uses internal coordinates instead of Cartesian coordinates and defines the potential energy as follows:

Potential energy 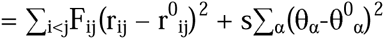, where

- r_ij_ is the distance between the atoms i and j, and the super-index 0 indicates the initial equilibrium conformation.
- F_ij_ is the matrix whose elements describe the force constant associated with each atom pair. F_ij_ = k/(1 + (r^0^_ij_/r_o_)^p^) if r^0^_ij_ < r_cut_, otherwise F_ij_ = 0 and k, r_o_, p and r_cut_ were set to 1, 3.8Å, 6 and 10Å respectively.
- The second term of the energy equation is added for tip effect prevention. θ_α_ is the dihedral angle [53].

Two applications, imode and imodview of iMOD were used in this study. First, imode program was used with –save_cart option to produce Cartesian normal modes in the output file with .evec extension. Second, the output file was used as an input file for imodview to compute the 3D vector sets of residues and ligands for the ten lowest frequency modes (n = 1 to 10). Residue/ligand molecule was set to be the averaging level to compute the arrow of eigenvector (level =1). According to recent studies, the first ten lowest frequency modes cover nearly 90% of protein conformational change, and thus, it is necessary for the training attributes consisting of information from the first 10 lowest frequency modes [25]. The NMA Training Attribute Value (NTAV) is defined as follows:

NMA Training Attribute Value = (DPVV_1_^2^ + … + DPVV_j_^2^ + … + DPVV_10_^2^)^1/2^, where j =1 to 10 is the normal mode index. DPVV is either the dot product of ligand displacement vector after normalization and residue displacement vector in DS-RMLR or the dot product of two displacement vectors of a residue upon ligand binding in DS-RMRR. Specifically, after aligning to the corresponding ligand-bound complex, closed-flap HIV-1 protease, 3IXO (PDB ID code), was used as the unbound structure of HIV-1 protease.

### Phase 4 Machine Learning

Machine learning is composed of a training phase and a predicting phase. Because two kinetic rate constants (k_off_, k_on_) are correlated, we used a multi-target random forest classification algorithm of Clus [54] to train coherent binary-output models to predict the two kinetic rate constants simultaneously.

- **Cross-Validation:** In this study, leave-one-out (LOO) cross-validation is used.
- **Performance Measurement:** Figure S12 depicts a confusion matrix for a binary classifier with two outcomes, high and low binding affinity (1,0). The targets that are correctly classified are denoted as true positives (TP) and true negatives (TN), and the targets that are misclassified are denoted as false positives (FP) and false negatives (FN). Sensitivity and specificity are the true positive (TP) and true negative (TN) rate respectively.

Common measures are accuracy and error rate. Accuracy is the percentage of a test set that is correctly classified and error rate is simply the percentage of a test set that is misclassified. They are computed as:

sensitivity = TP / (TP + FN)
specificity = TN / (TN + FP)
accuracy = 100 x (TP + TN) / (TP + FN + TN + FP) %
error rate = 100 x (FP + FN) / (TP + FN + TN + FP) %

Specifically, accuracy of the combined four-class log_10_k_on_/log_10_k_off_ binary prediction is defined as: if the true value is (1, 1), the accuracy for predicted (1, 1), (1, 0), (0, 1), and (0, 0) is 100, 50, 50, 0%, respectively.

- **Feature Selection:** Statistical experiment was conducted to identify the training features in DS-PIE, DS-EE, DS-RMLR, and DS-RMRR with frequency of occurrence greater than 25% in the LOO cross-validation experiment of the binary-target random forest classification algorithm.

---

**Table S1.** Results of the discretization.

| Binary Class | (0,0) | (0,1) | (1,0) | (1,1) |
| --- | --- | --- | --- | --- |
| Ligand I.D. | A037 | Saquinavir | B347 | B369 |
|  | B429 | B440 | B365 | B388 |
|  | B409 | Nelfinavir | A016 | A021 |
|  | A038 | Indinavir | A024 | B355 |
|  | B412 | B408 | A047 | A030 |
|  | B439 | Ritonavir | A023 | B322 |
|  | B268 | Amp | A017 | B425 |
|  | B277 | U75875 | B249 | A045 |
|  | B435 |  | A018 | B295 |
|  |  |  | A015 | B376 |
|  |  |  |  | A008 |
|  |  |  |  | DMP323 |

There are 9, 8, 10, and 12 training records in the binary classes of (0,0), (0,1), (1,0), and (1,1), respectively. Each record was identified by its corresponding ligand name.

**Table S2.**
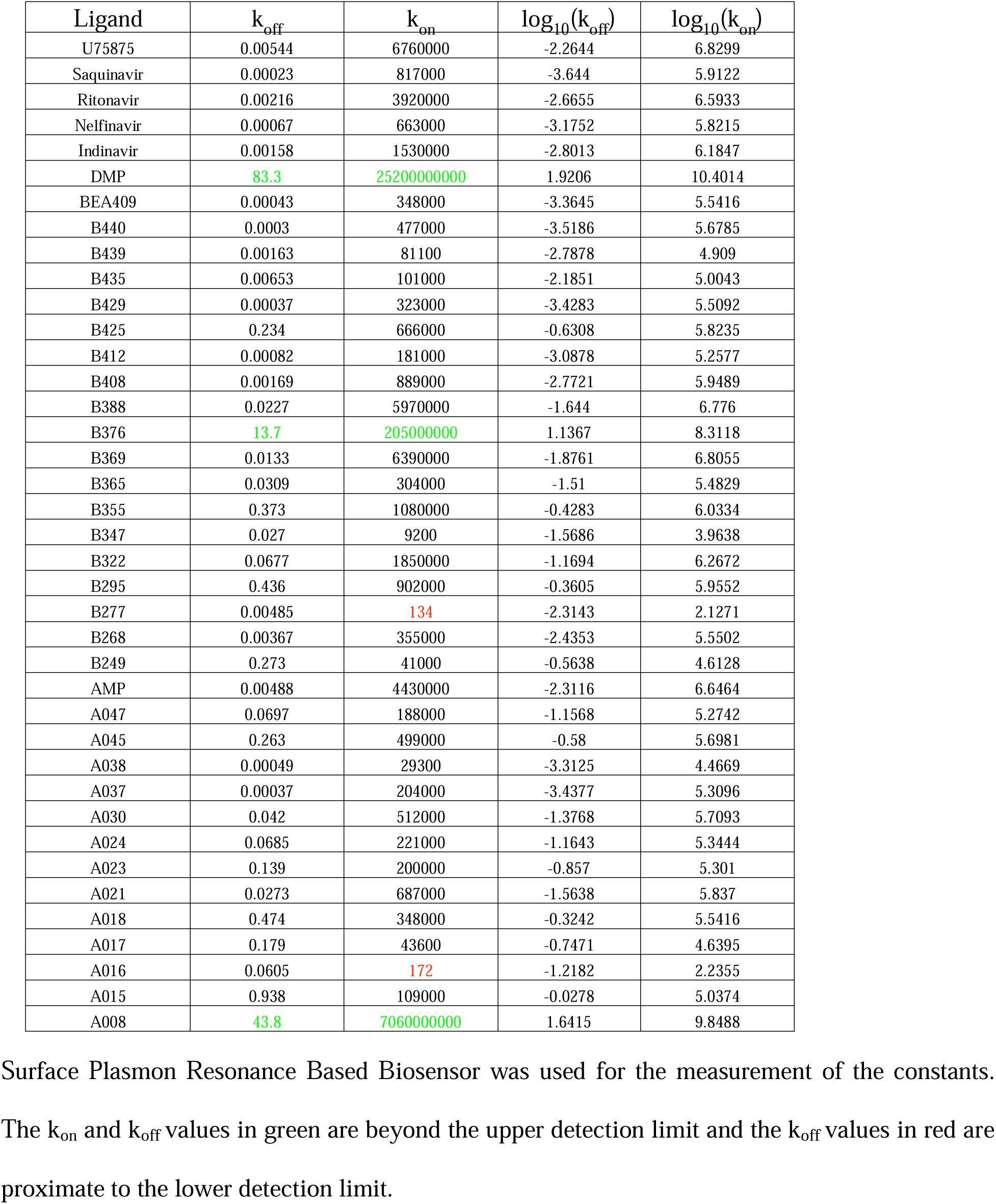
Association and dissociation rate constants (k_on_, k_off_) of the interactions between thirty-nine inhibitors and HIV-1 protease.

**Table S3.**
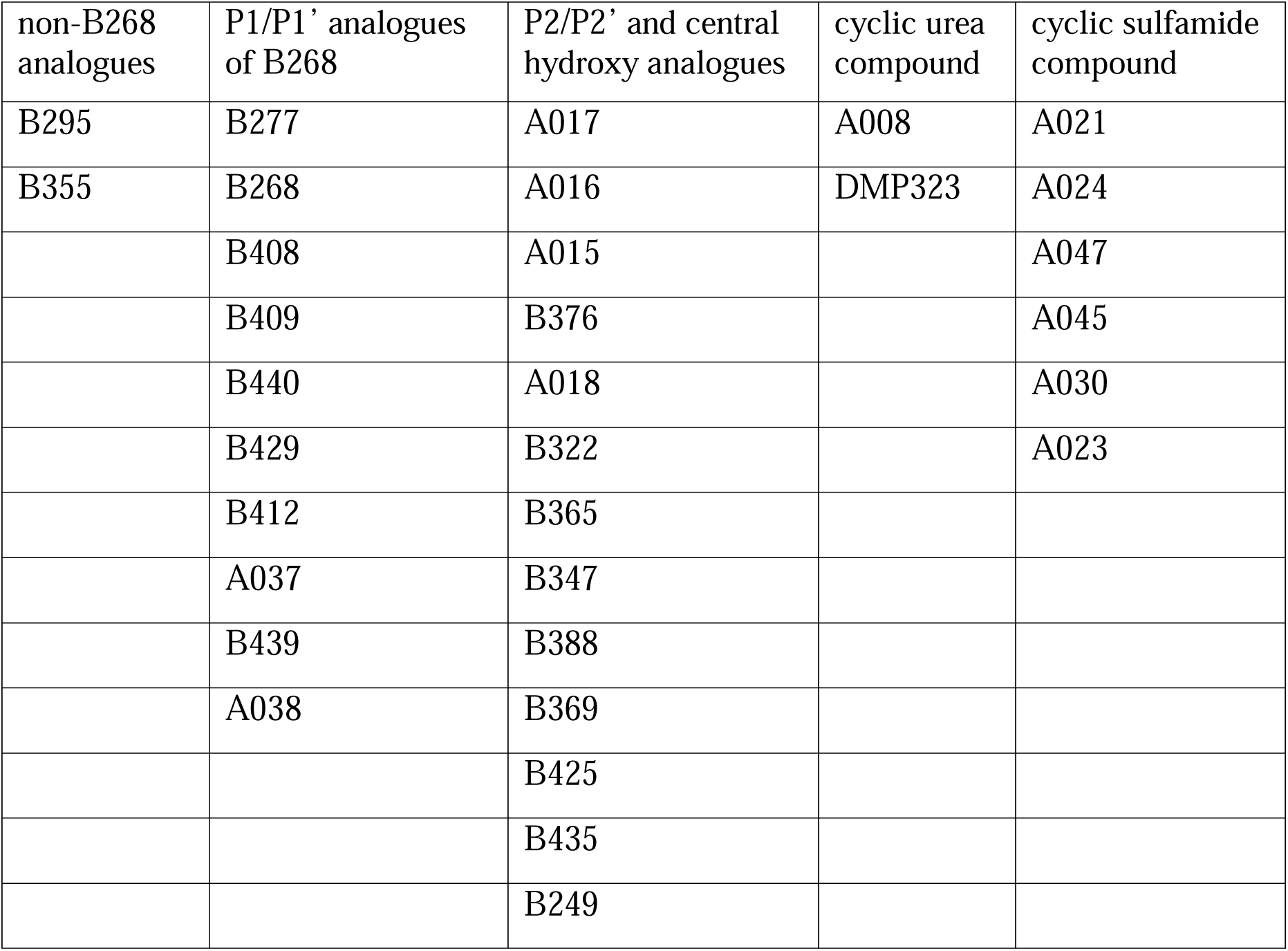
Structural classification of thirty-three ligand-bound HIV-1 complexes in reference to the structure of B268.

**Table S4.** eHiTS docking results.

| Complex | 1QBS | 1EBW | 1AJV | 1EC2 | 1D4H |
| --- | --- | --- | --- | --- | --- |
| Target Ligand | A008 | A015 | A021 | A037 | B295 |
|  |  | A016 | A023 | A038 | B355 |
|  |  | A017 | A024 | B412 | Indinavir |
|  |  | A018 | A030 | B429 | Saquinavir |
|  |  | B249 | A045 | B439 |  |
|  |  | B268 | A047 | B440 |  |
|  |  | B277 |  |  |  |
|  |  | B322 |  |  |  |
|  |  | B347 |  |  |  |
|  |  | B365 |  |  |  |
|  |  | B376 |  |  |  |
|  |  | B408 |  |  |  |

Target ligands and the HIV-1 protease of the complex in the same column were adopted in the eHiTS docking process to generate the corresponding ligand-bound HIV-1 complex structures.

**Figure S1.**
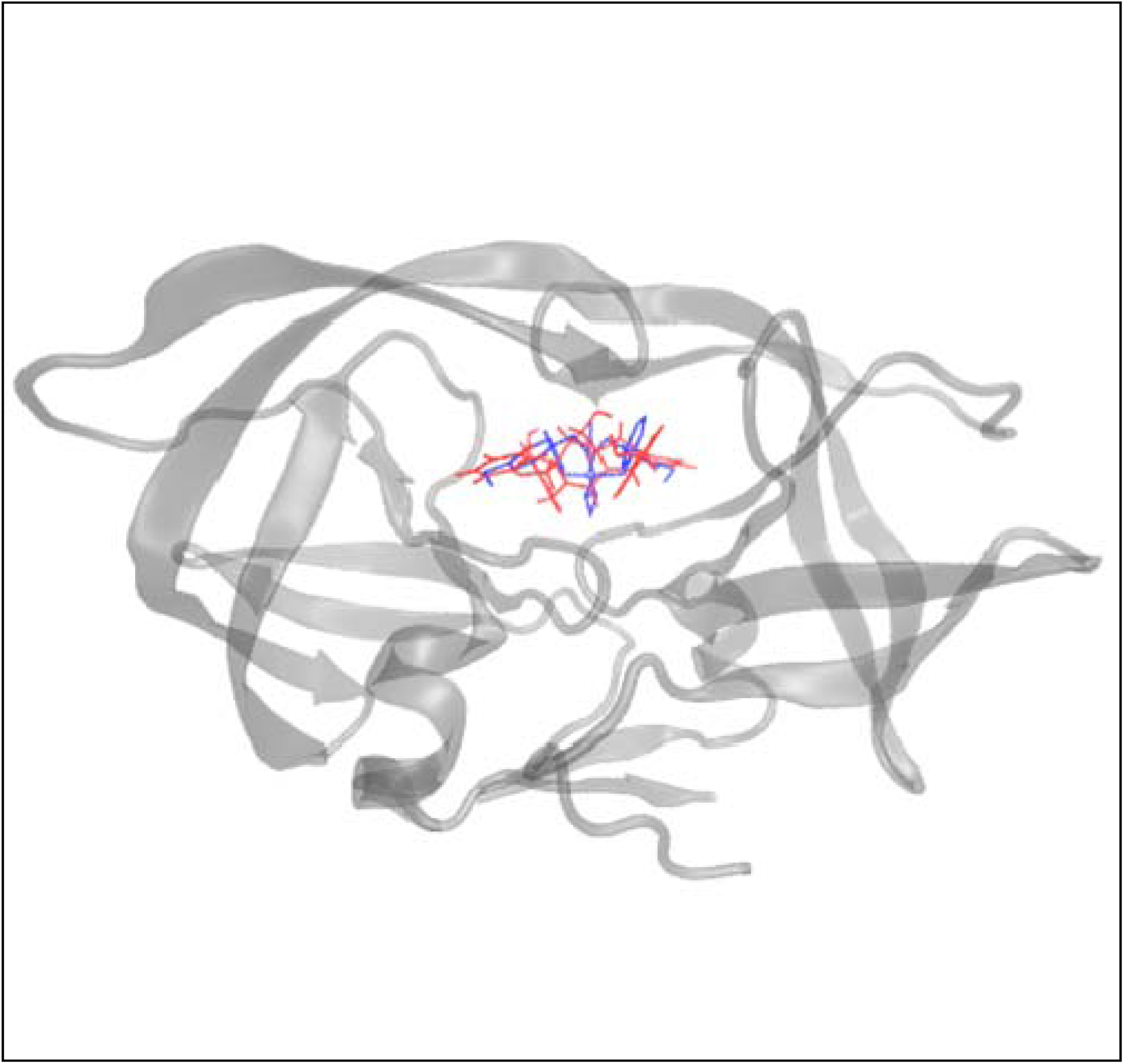
Docking ligand into HIV-1 protease. DMP (blue) is the co-crystalized ligand in HIV-1 protease (PDB code: 1QBS). Ligand A008 (red) is docked into the HIV-1 of 1QBS.

**Figure S2.**
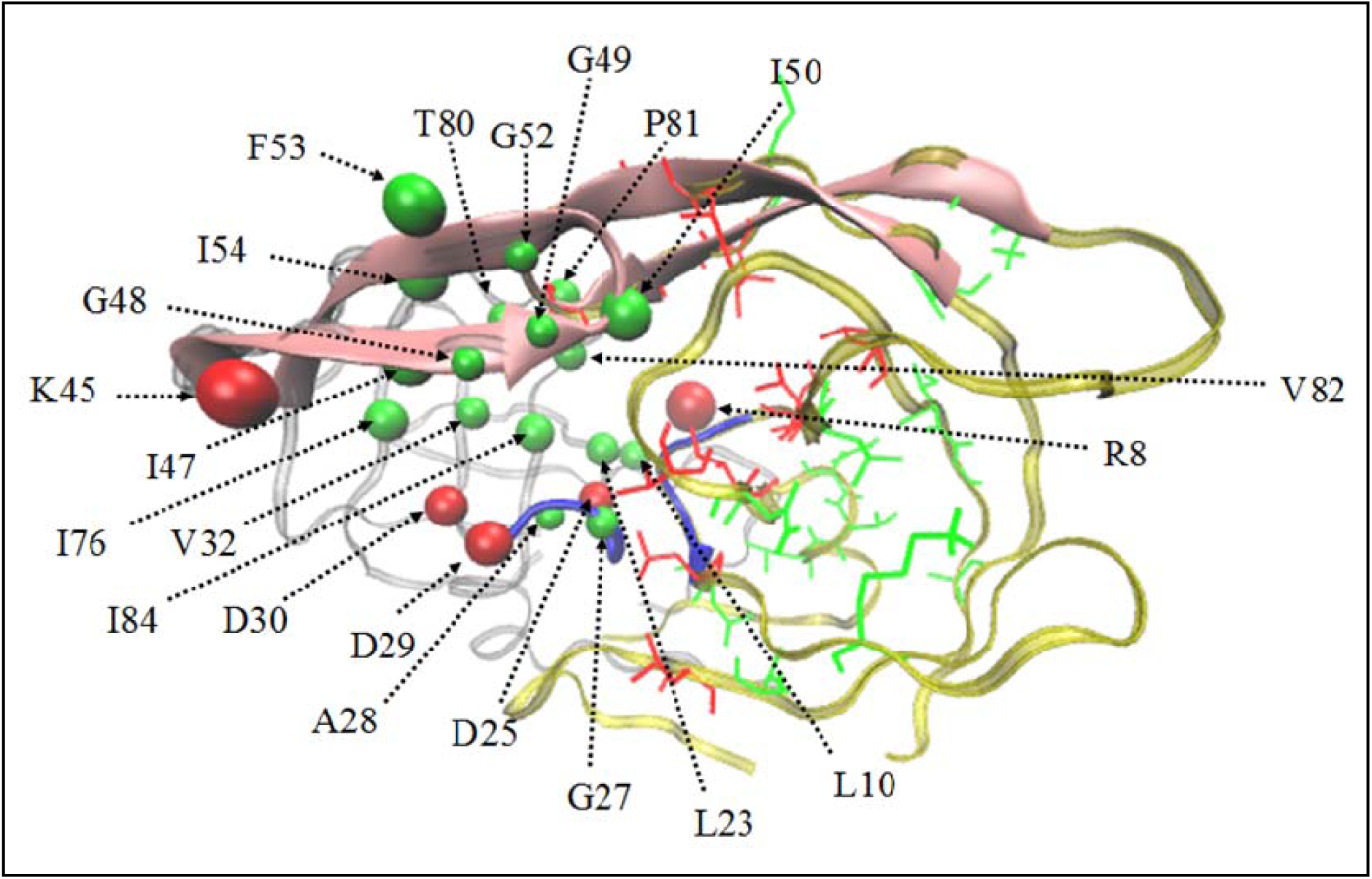
Forty-four HIV-1 residues selected by SASA program and the 26 drug resistant mutation residues. The chains A and B of HIV-1 are in transparent gray and green ribbons, respectively, with their flap regions (residue id: 43 – 58) in pink cartoon and active site (residue id: 25 – 29) in blue cartoon. The 22 SASA residues on the chain A are represented by 5 red beads for the charged residues and 17 green beads for the neutral residues. The 26 drug resistant mutation residues are depicted in lines on the chain B including the 12 PIRM residues near the binding site in red and the 14 PIRM residues outside the binding site in green.

**Figure S3.**
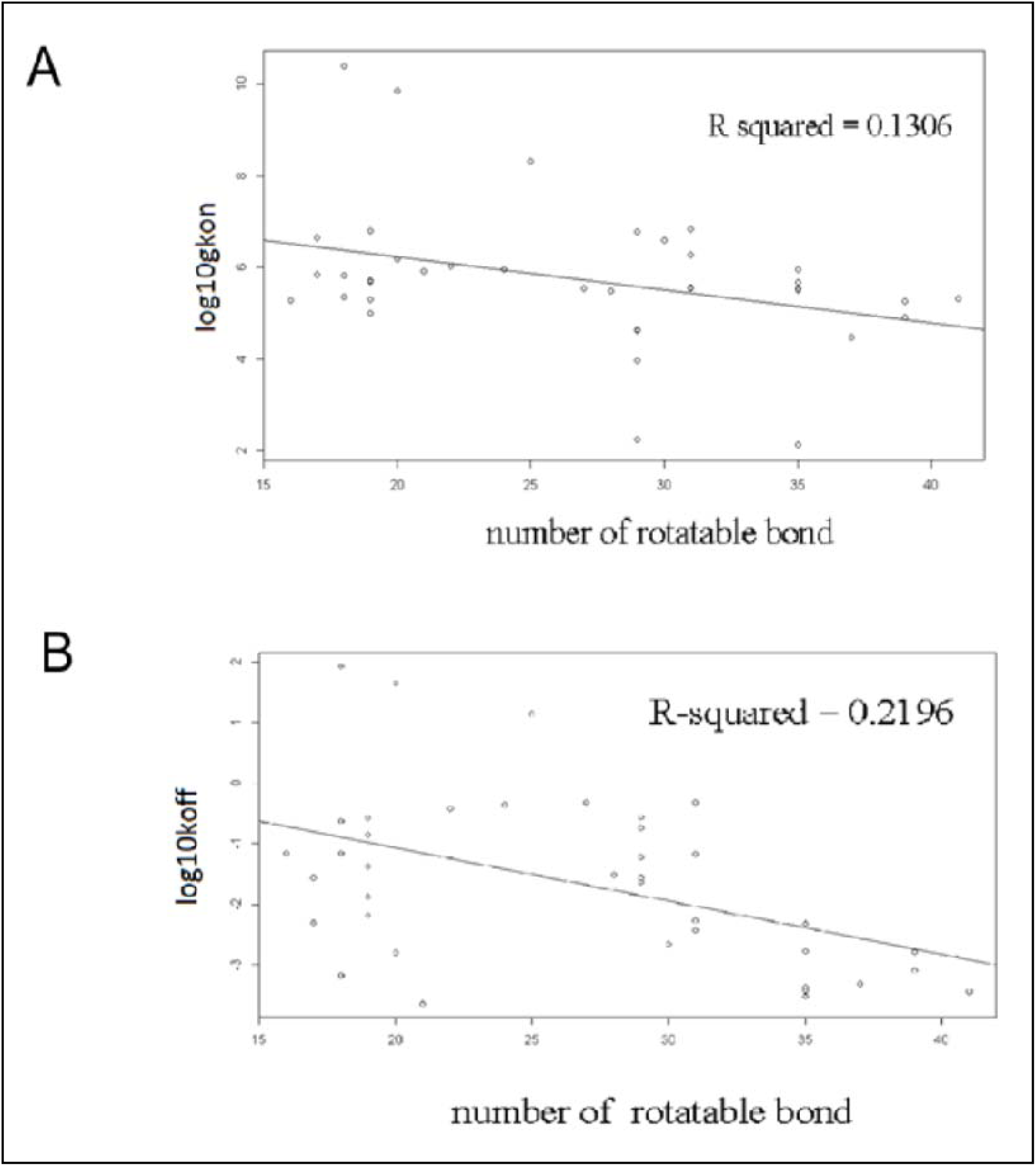
Number of ligand rotatable bond versus log_10_k_on_ / log_10_k_off_. Each data point represents one sample of ligand-HIV-1 complex.

**Figure S4.**
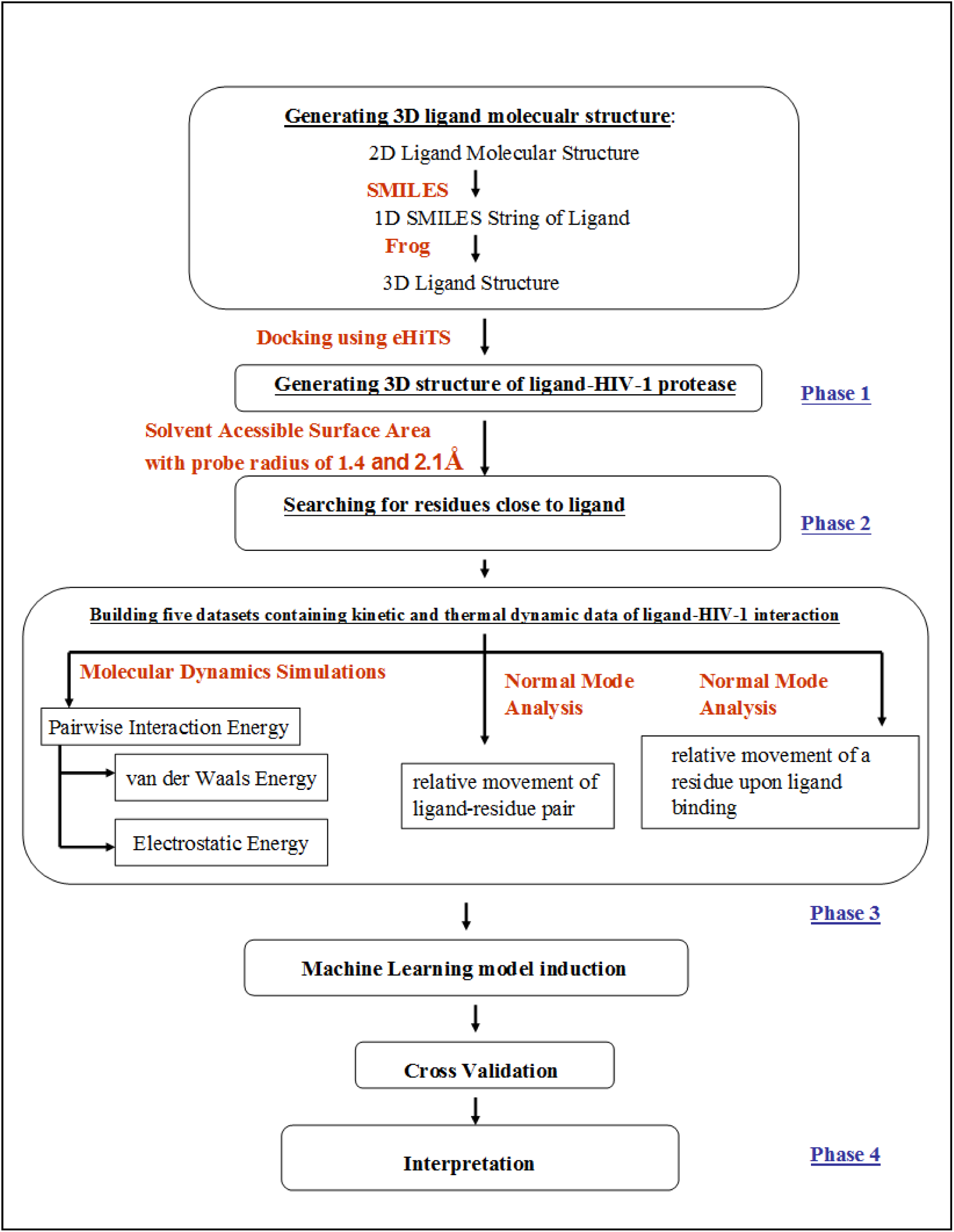
Schema of methodology.

**Figure S5.**
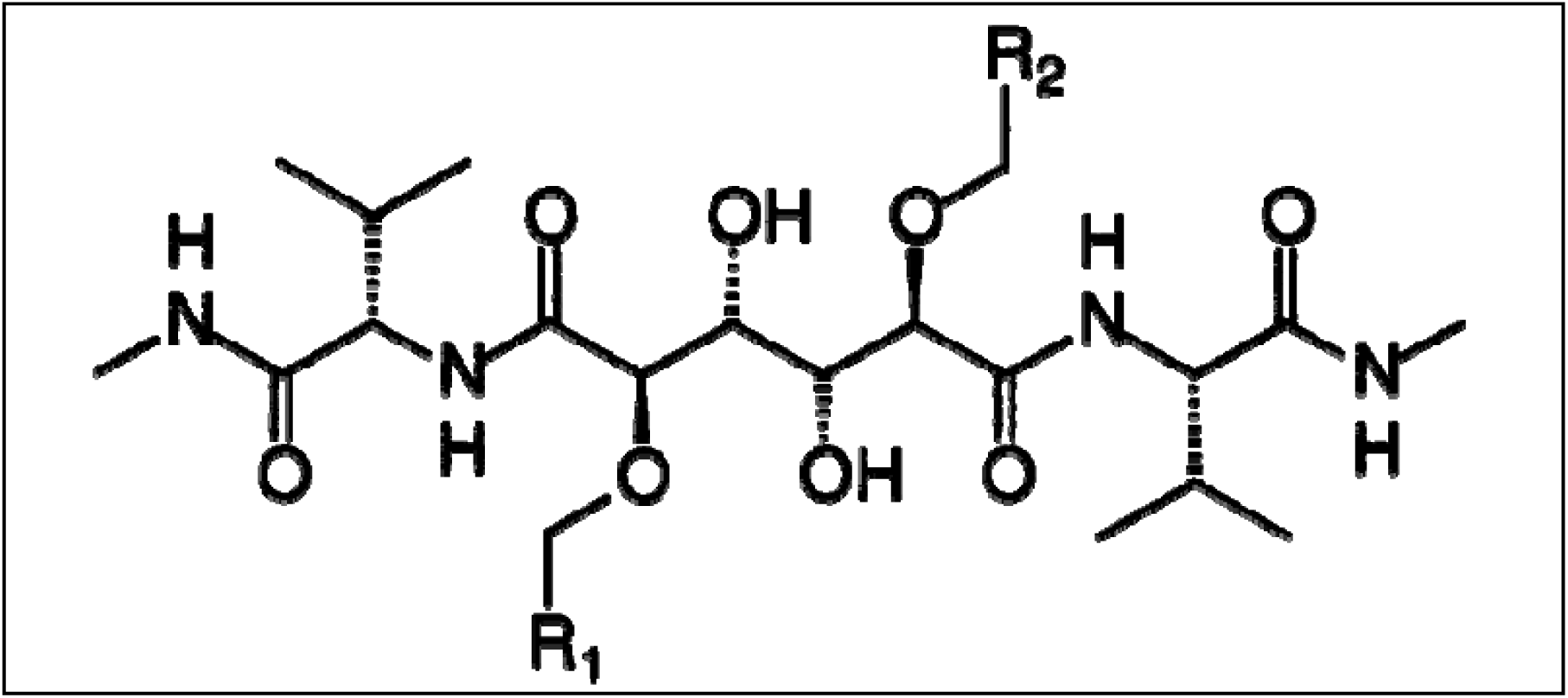
2D molecular structure of B268. R_1_ and R_2_ are benzene rings.

**Figure S6.**
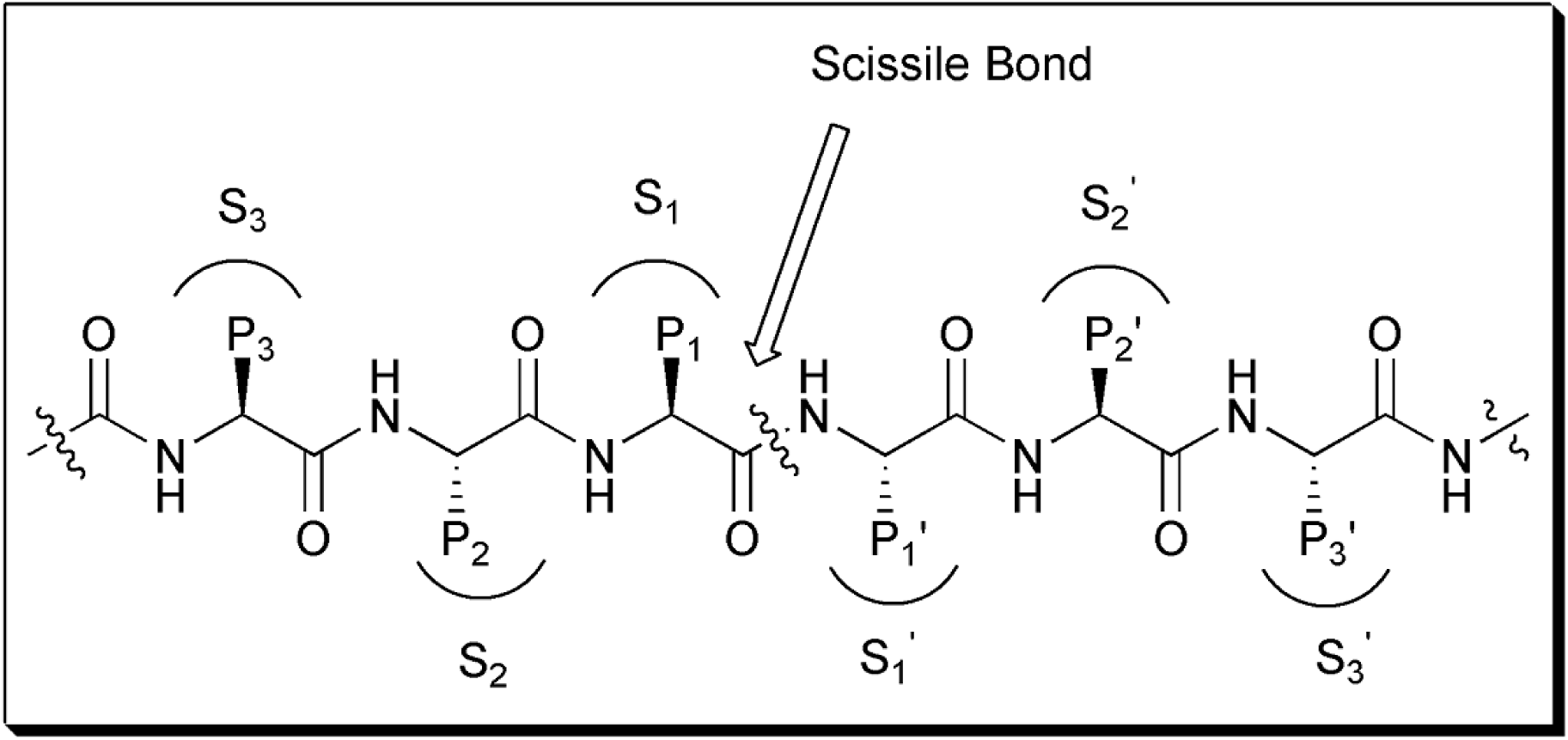
Structure of interaction between aspartyl protease and peptide substrate.

**Figure S7.**
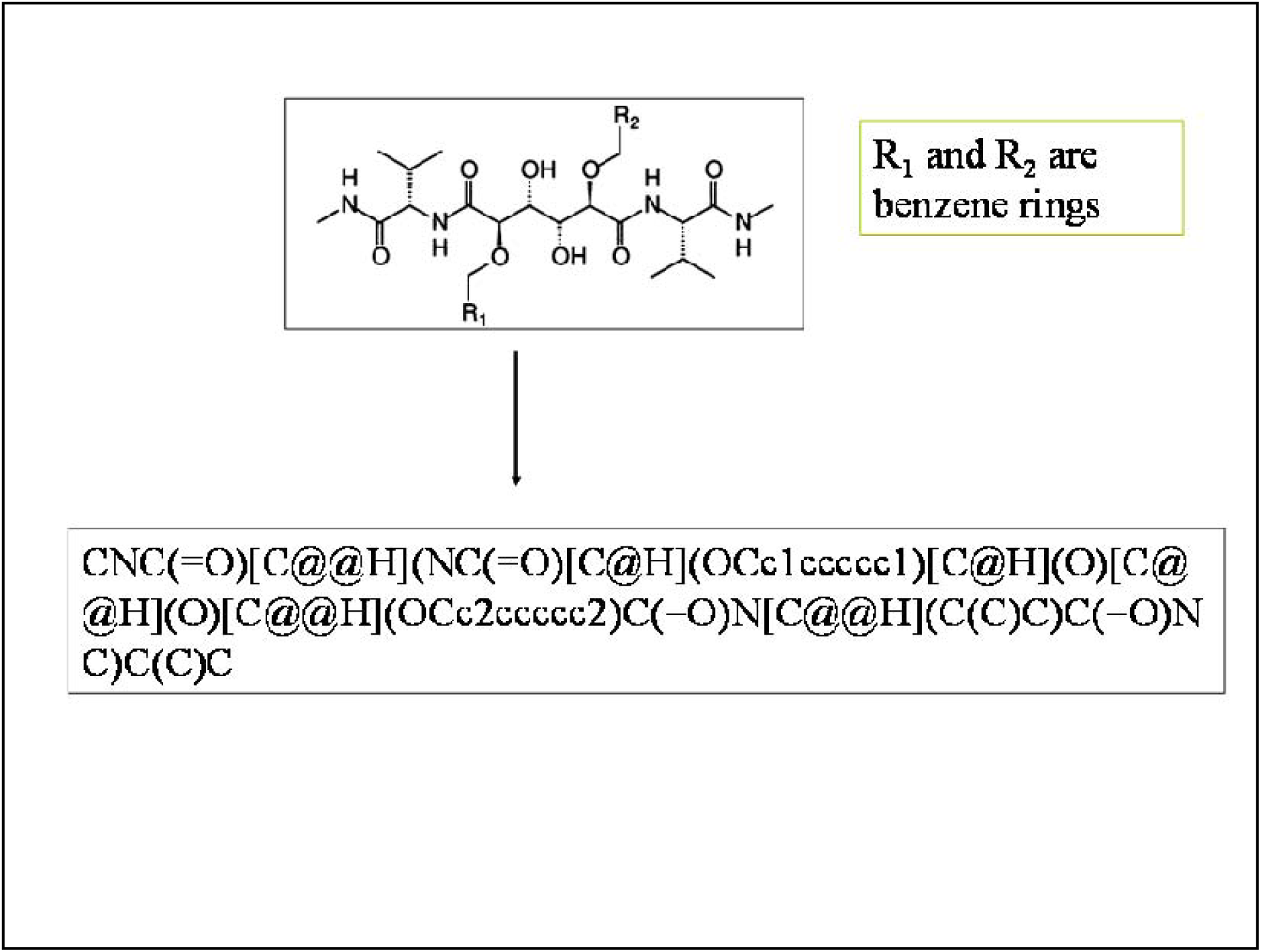
Transformation from the 2D molecular structure of B268 into its SMILES string.

**Figure S8.**
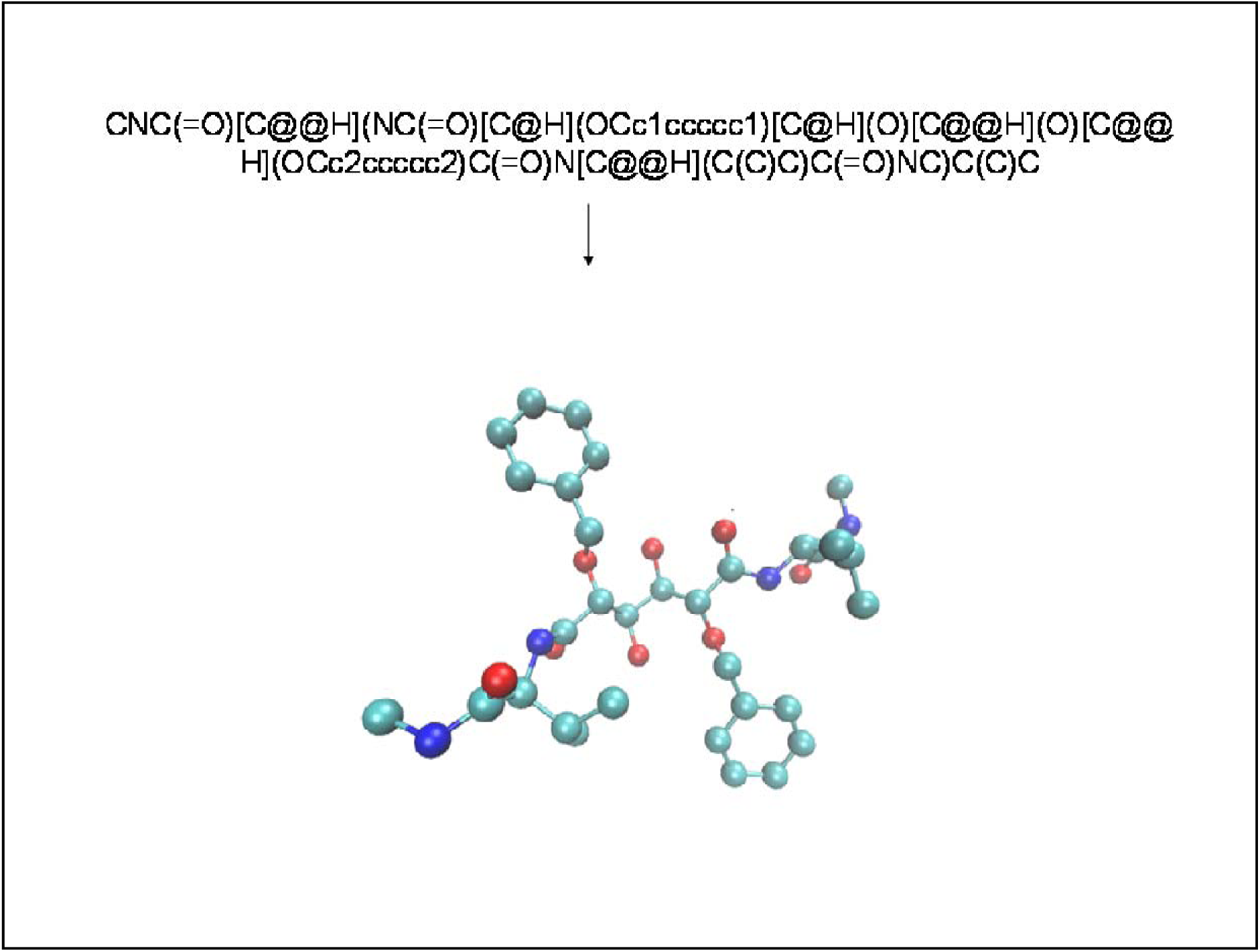
Conversion of B268 SMILES string into its 3D molecular structure. Color scheme: Turquoise, blue and red represent carbon, nitrogen and oxygen atoms respectively.

**Figure S9.**
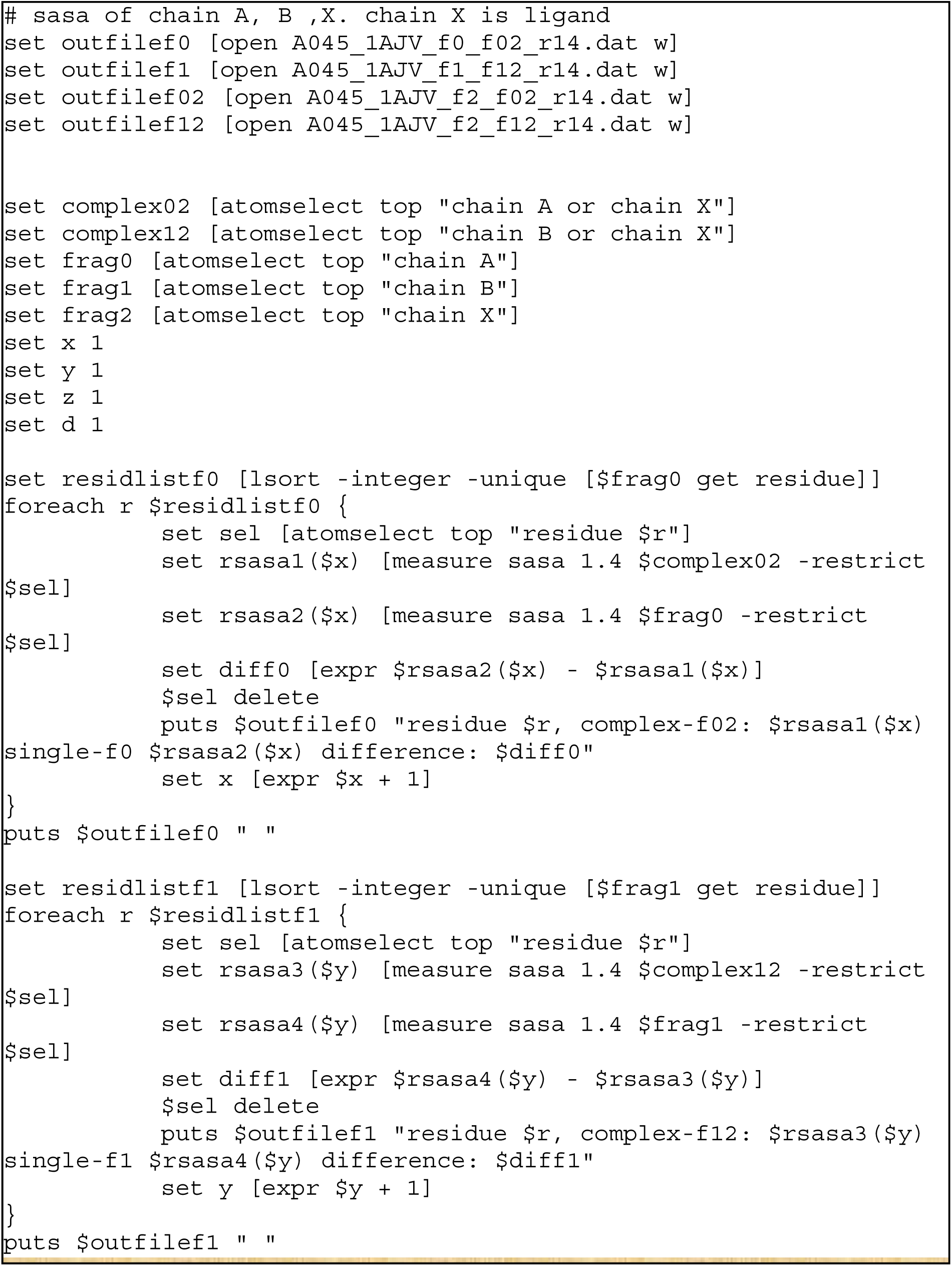
SASA TCL script with probe radius of 1.4A for the A045–1AJV complex.

**Figure S10.**
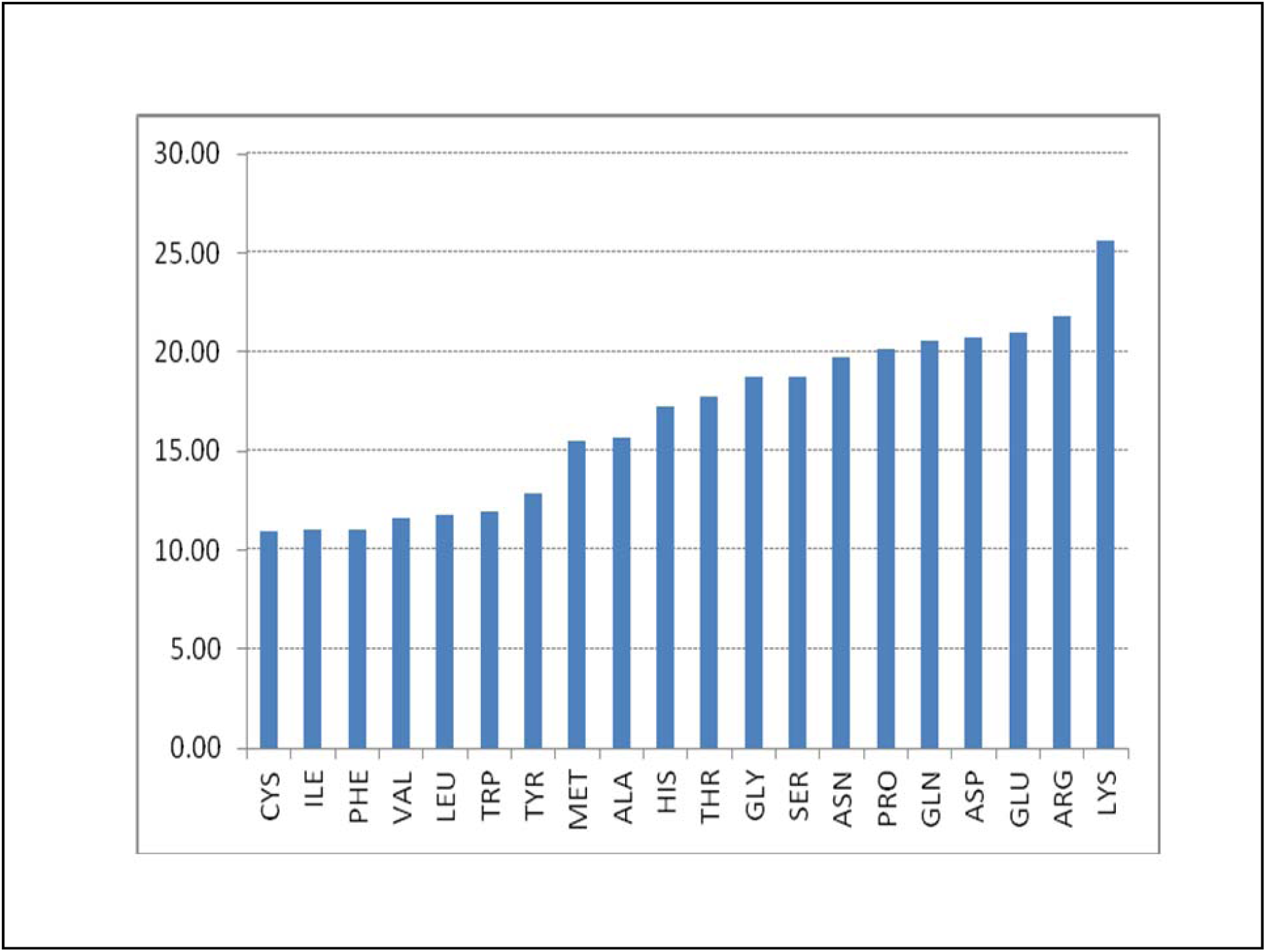
Average dielectric constants of different types of amino acids.

**Figure S11.**
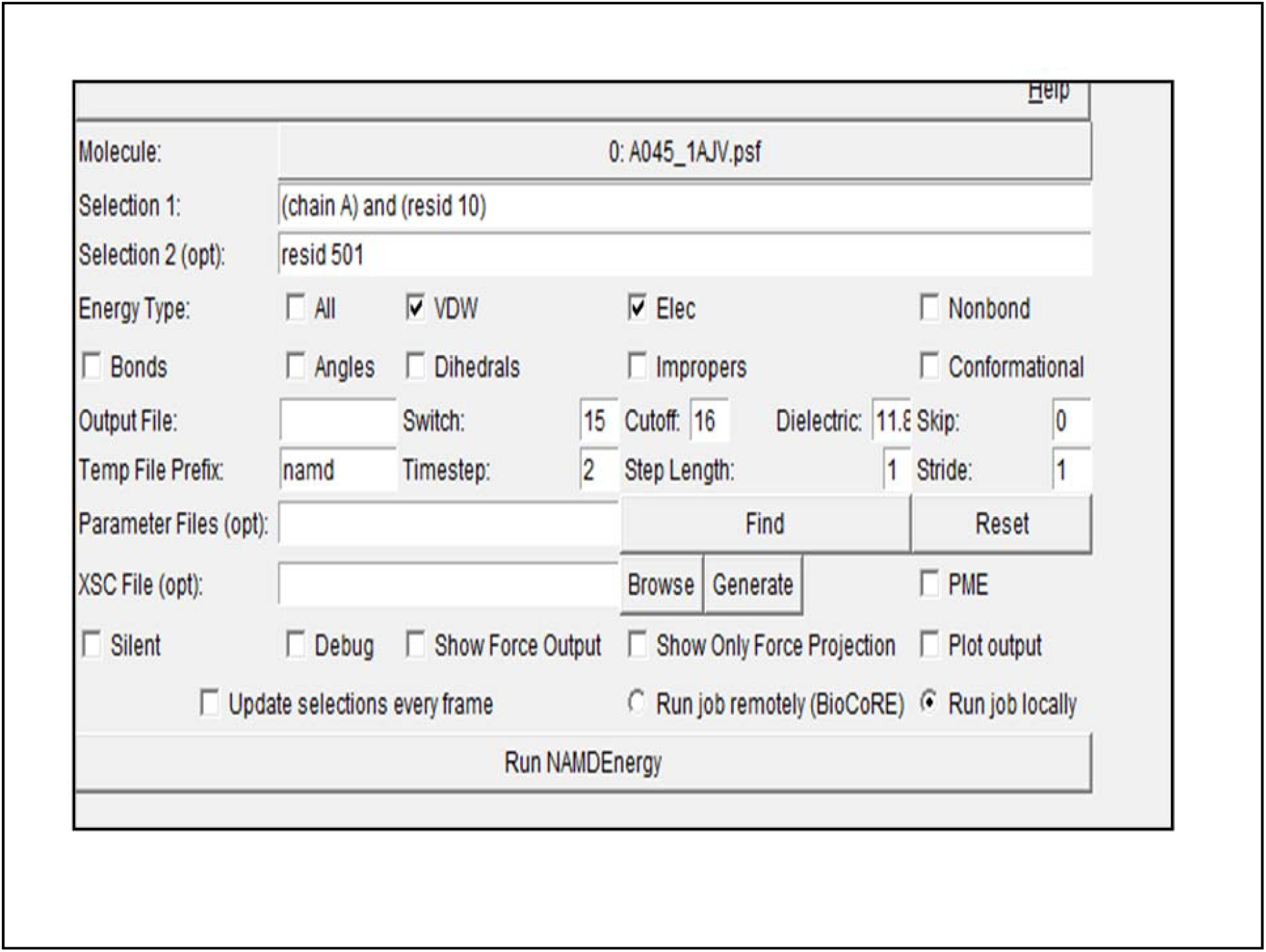
NAMDEnergy graphical user interface.

**Figure S12.**
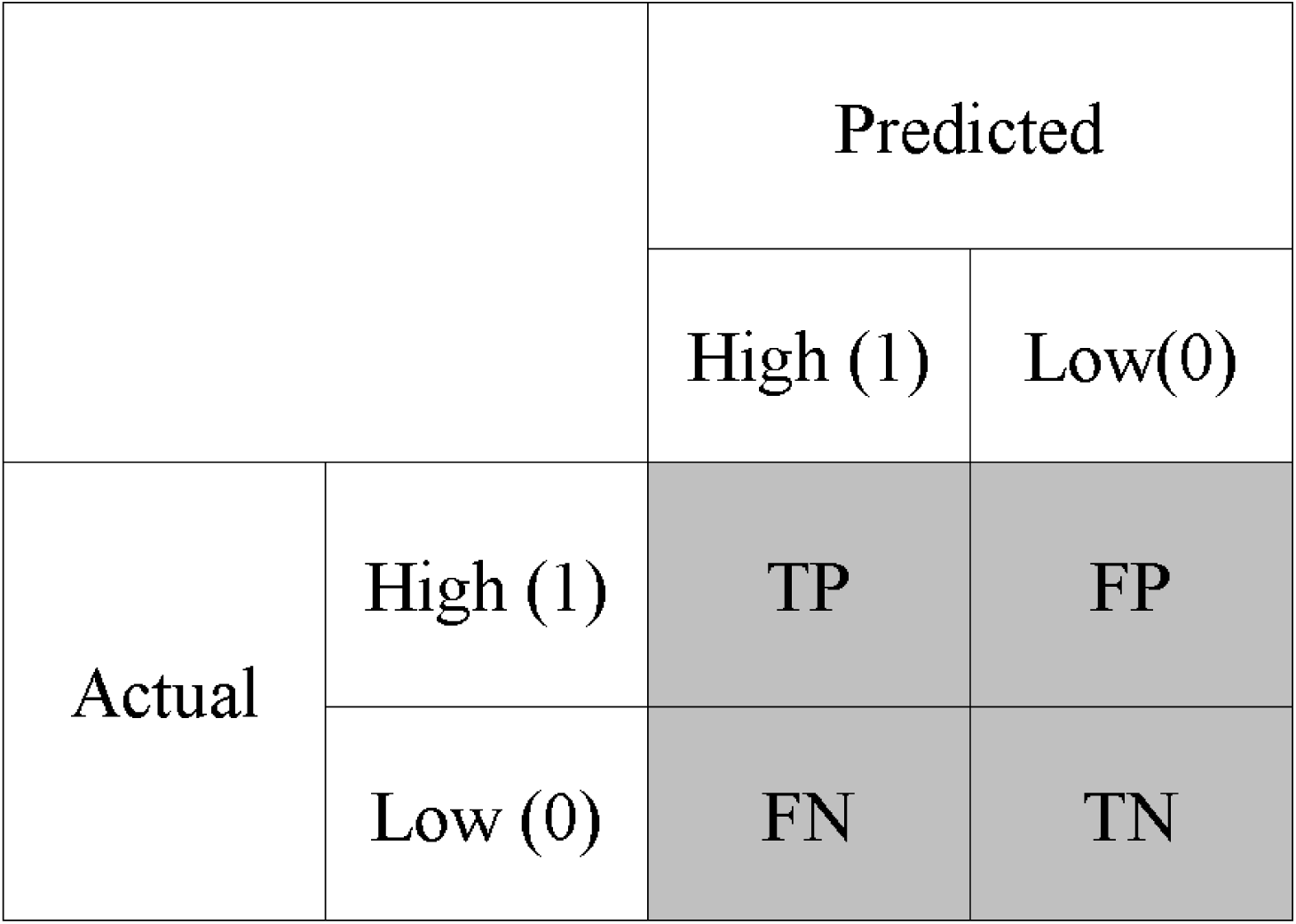
A confusion matrix for a binary classifier.

